# Context dependent activity of p63-bound gene regulatory elements

**DOI:** 10.1101/2024.05.09.593326

**Authors:** Abby A. McCann, Gabriele Baniulyte, Dana L. Woodstock, Morgan A. Sammons

## Abstract

The p53 family of transcription factors regulate numerous organismal processes including the development of skin and limbs, ciliogenesis, and preservation of genetic integrity and tumor suppression. p53 family members control these processes and gene expression networks through engagement with DNA sequences within gene regulatory elements. Whereas p53 binding to its cognate recognition sequence is strongly associated with transcriptional activation, p63 can mediate both activation and repression. How the DNA sequence of p63-bound gene regulatory elements is linked to these varied activities is not yet understood. Here, we use massively parallel reporter assays (MPRA) in a range of cellular and genetic contexts to investigate the influence of DNA sequence on p63-mediated transcription. Most regulatory elements with a p63 response element motif (p63RE) activate transcription, with those sites bound by p63 more frequently or adhering closer to canonical p53 family response element sequences driving higher transcriptional output. The most active regulatory elements are those also capable of binding p53. Elements uniquely bound by p63 have varied activity, with p63RE-mediated repression associated with lower overall GC content in flanking sequences. Comparison of activity across cell lines suggests differential activity of elements may be regulated by a combination of p63 abundance or context-specific cofactors. Finally, changes in p63 isoform expression dramatically alters regulatory element activity, primarily shifting inactive elements towards a strong p63-dependent activity. Our analysis of p63-bound gene regulatory elements provides new insight into how sequence, cellular context, and other transcription factors influence p63-dependent transcription. These studies provide a framework for understanding how p63 genomic binding locally regulates transcription. Additionally, these results can be extended to investigate the influence of sequence content, genomic context, chromatin structure on the interplay between p63 isoforms and p53 family paralogs.

## INTRODUCTION

Transcription factors regulate gene expression networks during development and are responsible for the maintenance of cellular and organismal homeostasis. These activities require transcription factor interactions with DNA, usually via conserved sequence motifs within cis-regulatory elements (CRE) like promoters and enhancers (Slattery et al., 2014). Sequence specific transcription factor binding to regulatory elements can affect gene expression in multiple, context-dependent ways, including direct recruitment of cofactors or RNA polymerase and through control of local and long-distance chromatin structure. TF control of gene expression is not a binary “on/off” state, and represents a range of dynamic interactions with DNA dictated by sequence, chromatin, and other locally-bound transcription factors (Hager et al., 2009; Ricci-Tam et al., 2021). Ultimately, understanding how DNA sequence and chromatin context at CREs controls TF binding is critical for dissecting complex gene regulatory networks during development and in disease.

The tight relationship between sequence-specific transcription factor activity, CREs, and gene expression is especially important for lineage specification during development (Spitz and Furlong, 2012; Long et al., 2016; Barral and Zaret, 2023). The transcription factor p63, a member of the well-known p53 family, is a key regulator of epithelial lineage specification and self-renewal (Senoo et al., 2007; Melino et al., 2015; Li et al., 2023). Extensive work using p63 loss-of-function mouse models demonstrates the essentiality of p63 for development of limbs, digits, and craniofacial structures (Yang et al., 1998; Mills et al., 1999). These phenotypes are consistent with those in humans, where p63 mutations cause multiple disorders rooted in epithelial cell dysfunction, including EEC (Ectrodactyly, Ectodermal Dysplasia and Cleft lip or Cleft lip and palate), Limb-Mammary Syndrome (LMS), Rapp-Hodgkin Syndrome (RMS), and ADULT syndrome (Celli et al., 1999; Amiel et al., 2001; McGrath et al., 2001, 2001; van Bokhoven et al., 2001; Bougeard et al., 2003). These epithelial-associated activities underlie the importance of p63 in multiple organ systems during development and in post-development contexts (Fletcher et al., 2011; Yallowitz et al., 2014; Richardson et al., 2017; Song et al., 2018). Organismal-level phenotypes in mouse models and human disorders are consistent with the indispensable role of p63 in the formation and maintenance of both the epidermis and epithelial-derived cells and tissues.

Mutations within p63-bound CREs are also directly linked to human developmental disorders, suggesting p63 regulation of CREs is required for development (Rahimov et al., 2008; Thomason et al., 2010; Lin-Shiao et al., 2019). While multiple human disorders are linked to mutations that reduce p63 function, p63 hyperactivity and gain-of-function contribute to post-developmental disorders like cancer. Overexpression of p63 drives tumorigenesis in squamous cell carcinomas (Ramsey et al., 2013; Saladi et al., 2017; Abraham et al., 2018), while genetic rearrangements in *TP63* lead to gain-of-function activities of p63 fusion proteins important for lymphoma progression (Saladi et al., 2017; Ng et al., 2018; Moses et al., 2019; Wu et al., 2023, p. 63).

Like other transcription factors, p63 activity requires direct binding to specific DNA motifs within CREs (Yang et al., 2006; Perez et al., 2007; Lambert et al., 2018). The *TP63* gene encodes multiple transcript and protein isoforms (Mills et al., 1999; Candi et al., 2006; Murray-Zmijewski et al., 2006; Sethi et al., 2015; Marshall et al., 2021; Osterburg and Dötsch, 2022), with the two most prominent being TAp63ɑ and ΔNp63ɑ generated from alternative promoter usage. TAp63ɑ is an obligate transcriptional activator, and functions in preservation of genetic integrity in germ cells, adult stem cell maintenance, and late-stage keratinocyte differentiation (Candi et al., 2006; Su et al., 2009; Gebel et al., 2017). On the other hand, ΔNp63ɑ’s activity is strongly context-dependent, and has been shown to be both a transcriptional activator and repressor (Fisher et al., 2020). ΔNp63ɑ is a pioneer factor which licenses epithelial-specific regulatory elements during development (Pattison et al., 2018; Li et al., 2019; Lin-Shiao et al., 2019; Yu et al., 2021), but can also bookmark chromatin structure at already established active regions or control 3D interactions to repress gene expression (Bao et al., 2015; Pattison et al., 2018). ΔNp63ɑ can also locally recruit traditional co-activators, like SMAD proteins and p300 (Krauskopf et al., 2018; Katoh et al., 2019; Klein et al., 2020, p. 300; Sundqvist et al., 2020), or co-repressors, like HDACs (LeBoeuf et al., 2010; Ramsey et al., 2011), to regulatory elements to variably control transcription (Sethi et al., 2014). The specific temporal and spatial contexts where ΔNp63ɑ performs these various transcriptional roles, and how these differential activities are regulated, remain unclear.

Gene regulatory elements contain multiple transcription factor binding sites in particular orientations that “code” for specific transcriptional outcomes. The combination of local sequence context and transcription factor occupancy at regulatory elements ultimately controls gene expression networks and vast cell fate decisions (Kulkarni and Arnosti, 2003; Zaret and Mango, 2016; Halfon, 2020). We implement STARR-seq MPRA technology to address whether sequence content and context of p63-bound gene regulatory elements might explain differential p63 activities like transcriptional activation or repression (Arnold et al., 2013). We identified sequence features within and around p63 binding sites that influence p63-specific activities. p63-mediated repression is most associated with local GC content and the presence of nearby motifs for known transcriptional repressors. p63-mediated activation is influenced by specific classes of p63 response element (RE) DNA motifs that also permit binding by p53. p63-bound CRE activity changes across different epithelial cell contexts with this variation potentially regulated by changes in p63 expression and flanking transcription factor motifs. ΔNp63ɑ occupancy is only weakly correlated to transcriptional output unlike strong transactivators like p53. However in the context of expression of a different isoform of p63, TAp63β, transcriptional activation greatly increases, suggesting that isoform switching is a mechanism that controls the activity of p63-bound regulatory elements.

## RESULTS

### Examination of the transcriptional regulatory potential of p63-bound elements

To explore how ΔNp63ɑ, hereby referred to as p63, controls epithelial gene regulatory networks, we measured transcriptional activity of putative cis-regulatory elements (CRE) bound by p63 using a massively parallel reporter assay (MPRA). We selected candidate CREs from a recent meta-analysis examining p63 ChIP-seq binding across multiple human epithelial-derived cell lines (Riege et al., 2020) and cloned them into a reporter system using the STARR-seq strategy (Muerdter et al., 2018; Neumayr et al., 2022)(Fig. 1A). We selected candidate CREs where p63 binding was observed in at least 8 independent ChIP-seq experiments, resulting in 17,310 elements. Sequences were cloned from single-stranded oligo pool where the likely p63 response element (p63RE) was placed in the center of the oligo, flanked by up to 52 nucleotides on either side of genomic context depending on the length of the identified p63RE. To understand the specific role of p63 in transcriptional regulation by the selected CREs, we created two variants predicted to disrupt p63 binding. We performed conservative substitutions of nucleotides enriched at any single position with greater than 80% frequency (p63RE mut) (Fig. 1B, Table S1). We also performed a random nucleotide shuffle of the predicted p63RE located at the center of the CRE as a control for disrupted p63 binding (p63RE shuffle). Finally, nucleotides flanking the p63RE (p63RE flank) or the entire CRE (full shuffle) were shuffled, all while preserving GC content of the original CRE sequence. The sequence of all variants can be found in Table S1.

**Figure 1.**
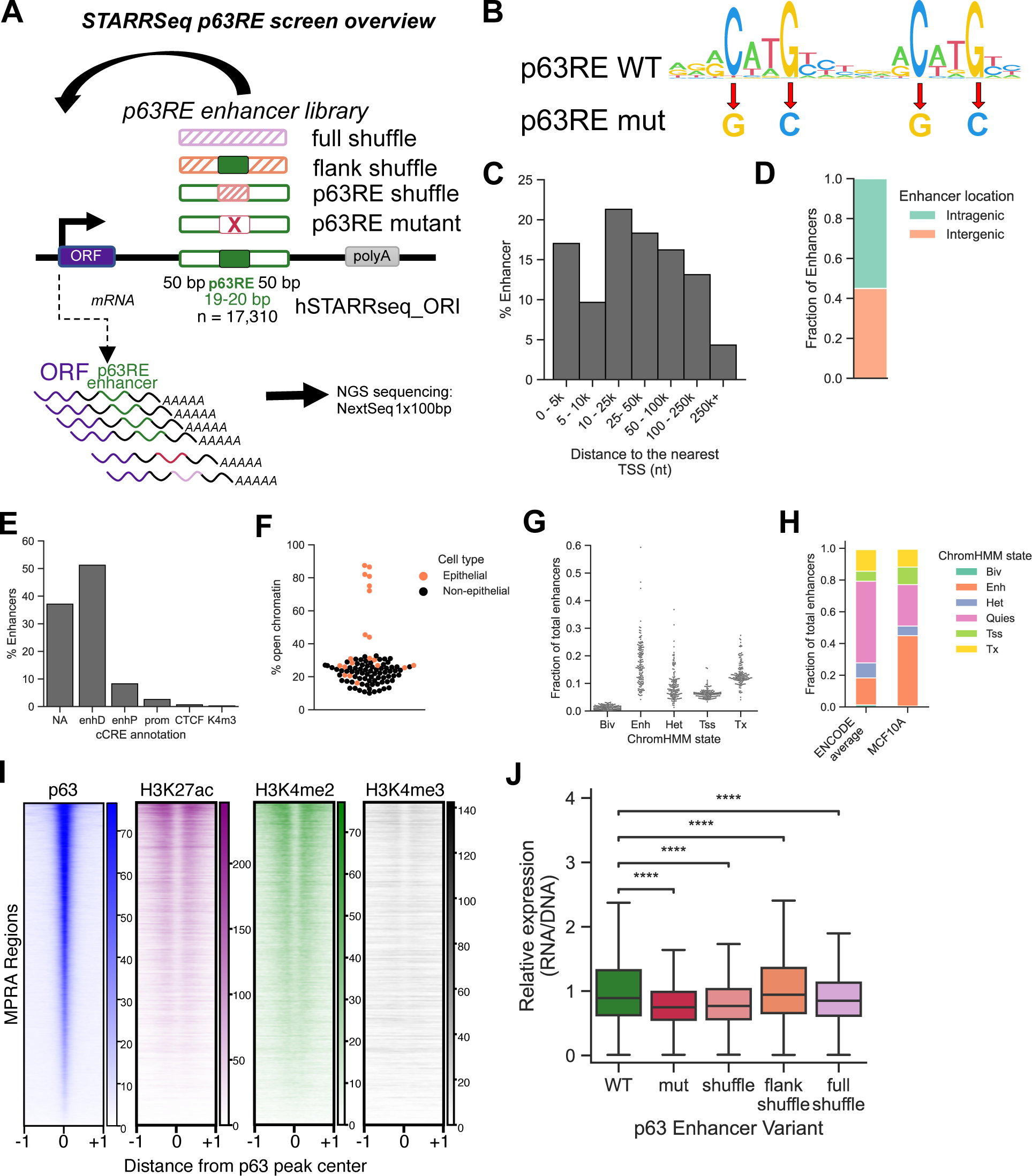
(A) STARR-Seq MPRA design. A p63 response element (RE) binding motif was centered within each putative cis-regulatory element (CRE) with a total length of up to 120 nucleotides. See Material and Methods and Table S1 for more information. Each wild-type (WT) element has a variant with p63RE mutation (mut or shuffle), scrambled flanking region (flankShuffle) or fully scrambled variant (fullShuffle). All elements preserve total GC content. (B) Position Weight Matrix (PWM) of the canonical WT p63RE or a mutant generated in this study (mut). (C) Distribution of CREs used in this study based on distance to the nearest RefSeq TSS. (D) Fraction of CREs found in intergenic or intragenic regions. (E) Distribution of CREs within ENCODE cCRE annotated functional elements. (F) Fraction of CREs occurring within DNAse Hypersensitive (DHS) clusters, denoted as “open”. G) Predicted regulatory status of CREs based on chromHMM chromatin state modeling. (I) Heatmaps representing p63, H3K27ac, H3K4me2 and H3K4me3 enrichment in MCF-10A cells at CRE locations used in this study. (J) Activity of CRE variants in MCF-10A cell line shown as a ratio of sequenced STARR-seq RNA reads to the original DNA library. *p*-values were calculated using Kruskal-Wallis test followed by Dunn’s *post-hoc* test and Bonferroni adjustment (\*\*\*\**p*-value < 0.0001).

We next examined the genomic and chromatin context of these elements to better understand how these characteristics might relate to their observed transcriptional output. Although many p63 binding events are intragenic (Fig. 1D), less than 20% of those are within 5kb of the transcriptional start site (TSS). Most sites are localized over 5kb from the TSS suggesting potential function as distal regulatory elements (Fig. 1C). ENCODE candidate Cis-Regulatory Element (cCRE) classification suggests that most of these p63-bound regions display distal enhancer-like signatures characterized by accessible chromatin and stereotypical histone modifications such as enrichment of H3K27ac and lack of H3K4me3 (Fig. 1E) (Moore et al., 2020). The next largest group overlaps the cCRE designation of NA which includes both heterochromatin and “quiescent” chromatin lacking known chromatin-based features of regulatory DNA. On average, 80% of these p63-bound elements are found in open chromatin regions of basal epithelial cell types (Fig. 1F) (Thurman et al., 2012; Sheffield et al., 2013). In contrast, most p63 binding sites are found in closed chromatin regions in all other cell types. We next examined the distribution of chromatin features surrounding our MPRA elements using chromHMM, which defines categorical chromatin states across multiple cell lines and conditions (Fig. 1G) (Ernst and Kellis, 2015; Vu and Ernst, 2022). In three p63-positive epithelial cell lines including the model mammary epithelial line MCF-10A, we observe enhancer-like enrichment at greater than 45% of the MPRA elements compared to an average of approximately 20% in non-epithelial cell lines (Fig. 1H). The epithelial specific enhancer-like chromatin features for the surveyed MPRA elements is consistent with epithelial lineage-restricted expression and pioneer factor activity of p63. We chose to use the model basal mammary epithelial cell line MCF-10A cell line for subsequent studies, as significant prior datasets for p63 occupancy, transcriptional regulation, and chromatin context are available. In line with summary statistics from ENCODE cCRE and chromHMM, the majority of our MPRA regions bound by p63 are enriched for H3K27ac and H3K4me2, but depleted for H3K4me3 (Fig. 1I), suggesting these elements have primarily enhancer-like qualities in MCF-10A.

We then assayed the activity of these p63-bound CRE to investigate sequence requirements for p63-dependent transcription in MCF-10A. Under basal conditions, these cells primarily express the p63 isoform ΔNp63ɑ, which is a context-dependent transcriptional activator or repressor (Fisher et al., 2020). We performed two biological replicates by transfecting the STARR-seq p63RE library into MCF-10A, isolated total RNA, and then specifically amplified and deep sequenced the self-transcribed CRE. Plasmid DNA pools were also sequenced as a transfection control, and CRE-driven RNA expression was quantified as a ratio of RNA:DNA. While 17,310 candidate p63-bound CREs were originally selected for analysis, synthesis, cloning, and experimental dropout reduced the number of regions used in downstream experiments and analysis to 13,696. We only included regions where all five variants were found in the DNA and RNA libraries at sufficient depth (Table S2, Materials and Methods).

First, we examined the role of the p63RE in mediating transcriptional activity. Mutation of either the entire p63RE (RE shuffle) or of specific nucleotides predicted to be critical for p63 binding (RE mut) substantially reduced transcriptional activation (p <= 1.00e-04) (Fig. 1J). Similar reductions in activity were seen when the entire CRE was shuffled, as expected by the loss of all native transcription factor binding sites. On the contrary, shuffling DNA sequence flanking the p63RE, and thus disrupting binding of other transcription factors, did not dramatically affect CRE-mediated transcription. These data suggest p63-bound CREs are more dependent on the central p63RE for transcriptional activation than other potential TF binding sites in flanking genomic context, similar to previous observations of the central importance for the p53 family RE at regulatory elements (Janky et al., 2014; Verfaillie et al., 2016; Sahu et al., 2022).

### Influence of p63 and p53 occupancy on cis-regulatory element activity

Given the importance of the central p63RE on CRE activity, we next investigated how p63 occupancy and enrichment at these elements influences transcriptional activity. We initially selected p63-bound regions for study that were identified in a meta-analysis of p63 ChIP-seq binding in between 8 and 20 independent ChIP-seq studies from different epithelial cell types, but did not contain p63 binding data from MCF-10A (Riege et al., 2020). We therefore used MCF-10A p63 ChIP-seq data from a prior study to examine how *in vivo* p63 enrichment was linked to CRE activity (Karsli Uzunbas et al., 2019). Increasing p63 enrichment in MCF-10A cells corresponds with increased p63 occupancy across epithelial cell types (Fig. 2A). In general, more ubiquitous p63 occupancy across cell types relates to p63 enrichment in MCF-10A, although significant variation exists across the range of binding events (Fig. 2A). CREs are more active when p63 binding occupancy is more ubiquitous compared to sites where p63 binding is restricted or cell-line dependent (Fig. 2B). On the contrary, MPRA activity is poorly correlated with p63 ChIP-seq enrichment in MCF-10A cells (Fig. 2C, Spearman ρ=0.127). These observations suggest that ubiquitous p63 binding, those events observed across many cell types, is more closely linked with transcriptional output than p63 binding enrichment as measured by ChIP-seq.

**Figure 2.**
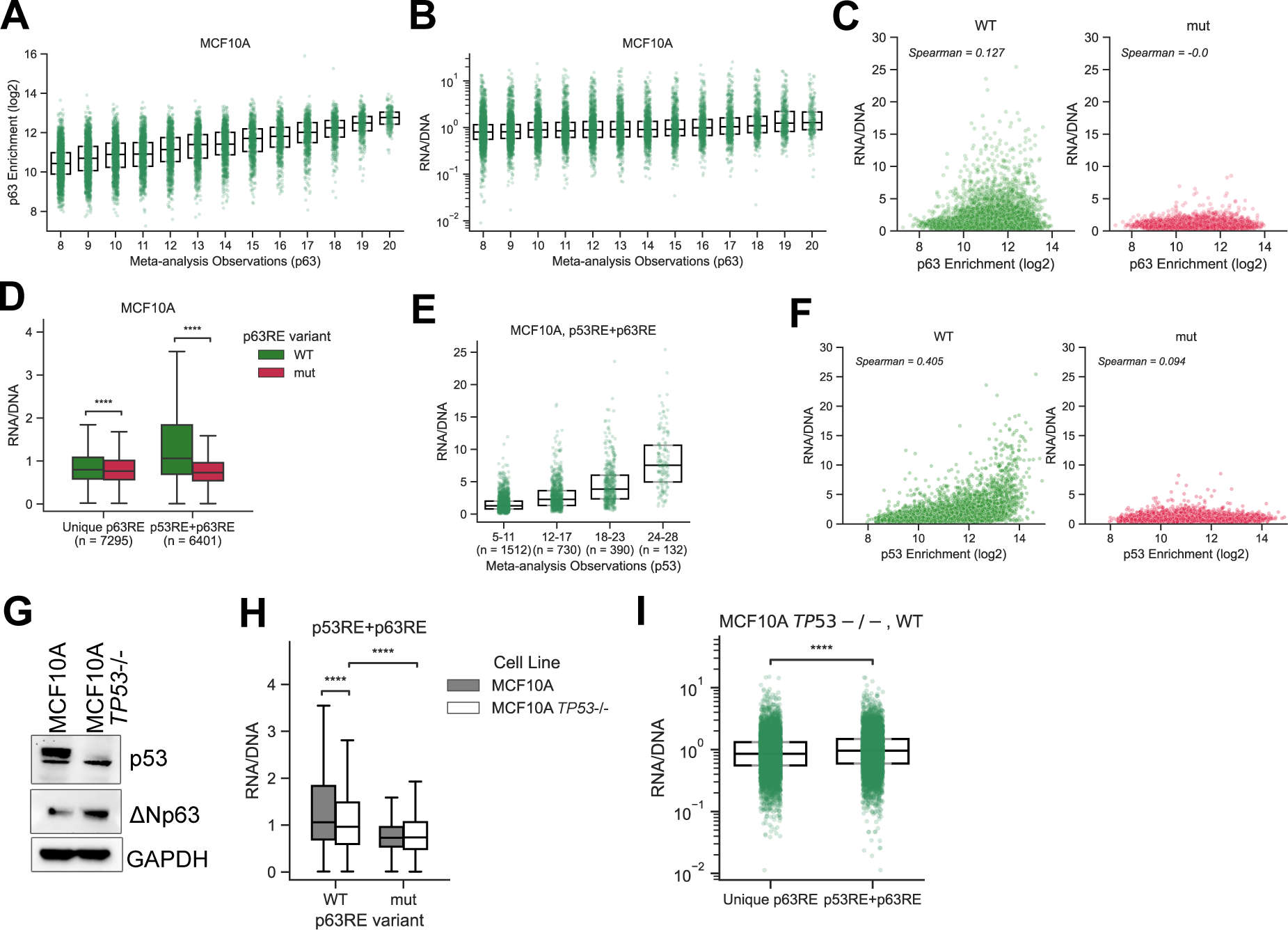
Box-plot showing relationship between meta-analysis-based p63 binding observation score and p63 ChIP-seq enrichment in the MCF-10A cell line (A) or WT CRE activity from the STARR-seq assay (B). (C) Correlation between WT or mut CRE activity and p63 ChIP-seq enrichment in MCF-10A cells (Spearman’s ρ=0.127, p=2.26e-48 for WT comparison and ρ=-0.0, p=0.98 for mut). (D) WT and mut CRE activity for Unique p63RE or p53RE+p63RE motif types with number of each motif type indicated on the *x*-axis (****: *p*-value < 0.0001, *Wilcoxon* signed-rank test) (E) Box-plot showing relationship between meta-analysis-based p53 observation score and p53 ChIP-seq enrichment in MCF-10A cell line. (F) Correlation between WT or mut activity and p53 ChIP-seq enrichment in MCF-10A cells (Spearman’s ρ=0.405, p=0.0 for WT comparison and ρ=-0.094, p=4.4e-27 for mut). (G) Western blot analysis of p53 and ΔNp63 expression in MCF-10A and MCF-10A TP53-/- cells. GAPDH is used as a loading control. (H) p53RE+p63RE enhancer activity in WT or TP53-/- MCF-10A cell lines (****: *p*-value < 0.0001, *Wilcoxon* signed-rank test). (I) Differences in WT CRE activity between p63RE motif types in MCF-10A TP53-/- cells.

Many p63 binding sites are shared by its family member p53, which is a near universal transactivator (Fischer et al., 2014; Sahu et al., 2022). Prior p63 ChIP-seq meta-analyses identified distinct p63 response element (RE) classes that differ primarily in their ability to support binding of p53 (p53RE+p63RE) or p63 only (unique p63RE) (Riege et al., 2020). Therefore, we examined whether intrinsic differences in p63RE motifs and overlapping binding with p53 might better reflect the observed transcriptional activity. CREs containing the p53RE+p63RE motif type were significantly more active than those with a unique p63RE (Fig. 2D), and saw a greater drop in activity when the central motif was lost. Similar to p63, p53 binding observations correlate with transcriptional output (Fig. 2B,E). Unlike our observations with p63, p53 binding strength from MCF-10A cells is better correlated with CRE activity (Fig. 2F, Spearman’s ρ = 0.405).

We next measured MPRA activity in MCF-10A *TP53* ^-/-^ cells (Fig. 2G) to parse specific contributions of p53 versus p63. CRE activity is significantly reduced in *TP53* ^-/-^ relative to WT MCF-10A cells (Fig. 2H). However, we also observe an additional significant reduction in activity when the central p63RE is mutated in *TP53* ^-/-^ conditions. Notably, in aggregate, p53RE+p63RE elements are more active than unique p63 CREs even in the absence of p53 (Fig. 2I). These data suggest that although p53 drives a substantial proportion of transcriptional activity for these CREs, p63 still functions as an activator at p53RE+p63RE motifs even in p53s absence. They also suggest that inherent sequence differences may contribute to differential transcriptional output of p63-bound CREs.

### Intrinsic response element sequence differences and GC content contribute to p63 and p53-dependent transcriptional activity

Initial analysis suggests that p63 primarily functions as a transcriptional activator, with mutations to the p63RE significantly reducing transcriptional output (Fig. 1J) despite p63 enrichment being poorly correlated with transcriptional activity (Fig. 2C). CREs containing p53RE+p63RE sequences are substantially more active than those containing motifs supporting only p63 binding (Fig. 2D) independent of whether p53 or p63 is engaged (Fig. 2I). The extent to which variation in half-site sequence and other intrinsic DNA information within a p53 family response element leads to differential binding kinetics and activities remains an open question in the field (Szak et al., 2001; Safieh et al., 2023).

p53RE+p63RE motifs were originally subdivided into five categories based on differences in occupancy and abundance in p53/p63 ChIP-seq datasets: primary, secondary, tertiary, quaternary, and quinary (Fig. 3A) (Riege et al., 2020). Primary motifs are considered the canonical p53 family motif, containing two canonical CWWG half-sites separated by a 6bp spacer (el-Deiry et al., 1992; Castro-Mondragon et al., 2022). CREs containing these elements support higher enrichment of p63 (Fig. 3B) and p53 (Fig. 3C) and are more active (Fig. 3D) compared to the other classes. The activity of secondary, quaternary, and quinary elements descend in that order (Fig. 3D), as do p63 and p53 occupancy (Fig. 3B,C). CRE activity significantly decreases when nucleotides critical for p53/p63 binding are mutated in both primary and secondary motifs (Fig. 3D). Because we selected CRE sequences based on p63 occupancy alone, our assays ultimately did not contain any tertiary p53RE+p63RE motifs. Likely due to their limited number in this dataset, mutation of quaternary (n=50) and quinary (n=82) motifs lead to a small, but not statistically significant, decrease in CRE activity (Fig. 3D). Similar to p53, p63 activates both primary and secondary motif-containing elements, but without a strong preference for primary motifs (Fig. 3E).

**Figure 3.**
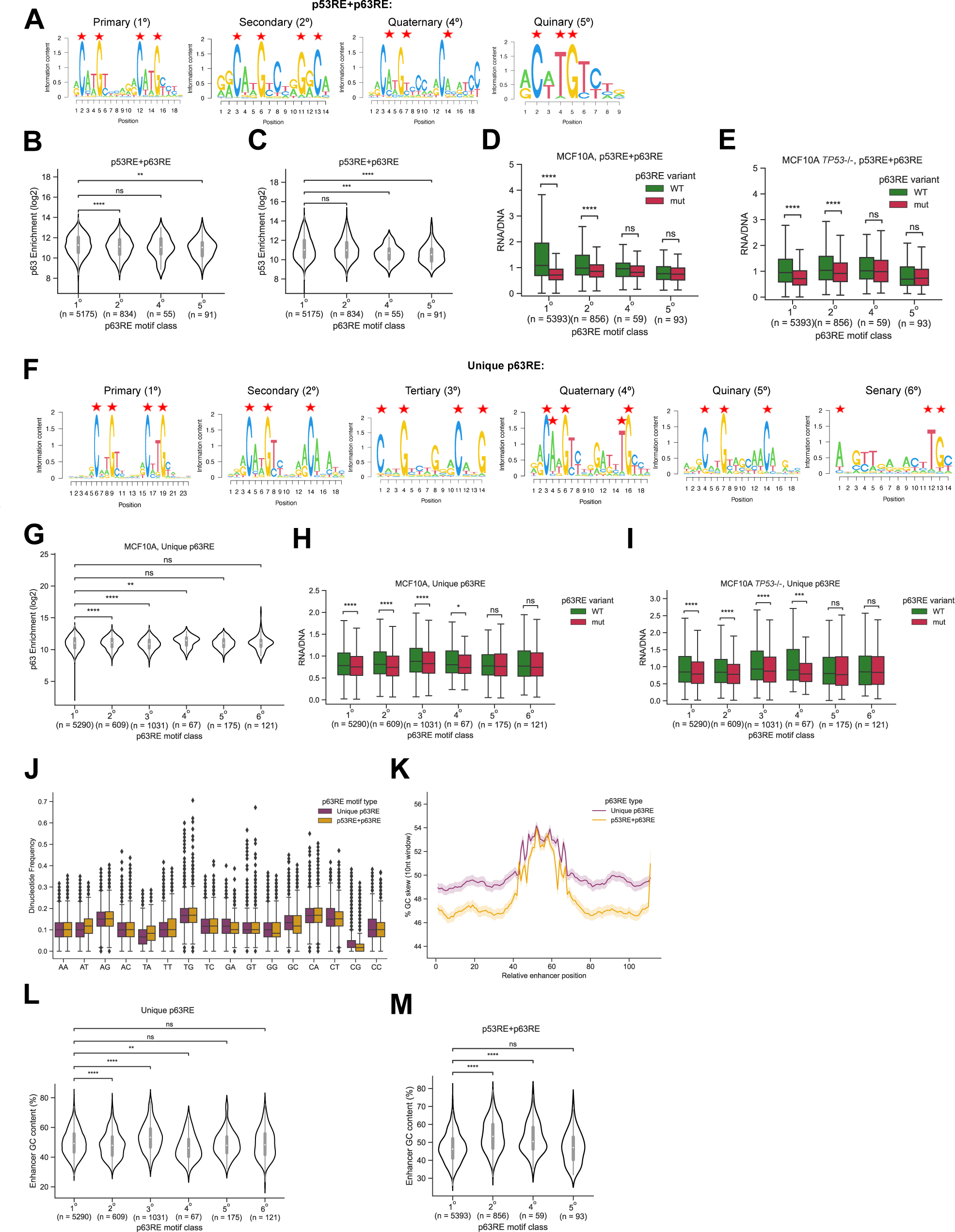
Analysis of p63RE motif class and CRE activity. (A) PWM of p53RE+p63RE motif classes, with stars indicating nucleotide substitutions in mut variants. p63 (B) or p53 (C) ChIP-seq enrichment in MCF-10A cells within each p53RE+p63RE motif class. (D) WT or mut enhancer activity in MCF-10A (D) or MCF-10A TP53-/- (E) within each p53RE+p63RE motif class. (F) PWM of Unique p63RE motif classes, with stars indicating nucleotide substitutions in mut variants. (G) p63 ChIP-seq enrichment in MCF-10A cells within each Unique p63RE motif class. WT or mut enhancer activity in MCF-10A (H) or MCF-10A TP53-/- (I) based on Unique p63RE motif class. (J) Dinucleotide repeat motif frequency in CREs. (K) Average GC content across CRE regions separated by motif type. GC content was determined using a 10 nt sliding window approach. Shaded area represents a 95% confidence interval. (L) Average CRE GC content of Unique p63RE (L) or p53RE+p63RE (M) within each motif class. Statistical comparisons were computed using either Mann-Whitney U test (B, C, G, L, M) or Wilcoxon signed-rank test (D, E, H, I). *p*-values are indicated as *ns*: 0.05 < *p*, **: 0.001 < *p* <= 0.01, ***: 0.0001 < *p* <= 0.001, ****: *p* <= 0.0001.

We next examined whether CRE activity might relate to specific classes of the unique p63RE motifs (Fig. 3F). The primary and tertiary motifs resemble canonical p53 family motifs, with two half sites separated by a 6bp spacer. Secondary, quaternary, and quinary motifs are characterized by the presence of a single half-site coupled with an incomplete second half-site. Senary motifs, relatively lowly represented in the test sequences, generally contain a single, weak half-site. Septenary motifs were excluded from this analysis due to low representation (n=2). The relationship between unique p63RE motif type and CRE activity is more nuanced than for those with p53RE+p63RE motifs. p63 enrichment is highest at primary and quaternary elements (Fig. 3G), but tertiary elements drive the highest level of CRE expression (Fig. 3H,I). Loss of the central element leads to statistically significant loss in CRE activity for primary, secondary, tertiary, and quaternary elements, with quinary and senary motifs driving the lowest expression and having the least dependence on the central motif. These observations are similar in the presence and absence of p53 consistent with lack of p53 binding at these sites (Fig. 3I)

Increasing p63 ChIP-seq enrichment is not coupled to increased activity (Fig. 2C), and different p63RE classes contribute to, but do not explain, differential CRE activity. We sought to determine whether other local sequence features might provide additional insight into p63-dependent CRE activity. Previous studies demonstrate dinucleotide repeat motifs (DRMs) are an indicator of regulatory sequence activity (Yanez-Cuna et al., 2012; White et al., 2013; Colbran et al., 2017). Dinucleotide content was generally similar within CRE classes with the exception of increased GC and CG DRM enrichment in unique p63RE and increased AT and TA enrichment for p53RE+p63RE (Fig. 3J). Increased AT and TA dinucleotide content in the p53RE+p63RE likely reflects the strong preference of p53 for these dinucleotides in half-sites (Fig. 3A, 3J). CG dinucleotide enrichment and subsequent methylation could potentially explain reduced activity of unique p63RE compared to p53RE+p63RE, but these observations require further study.

Differences in the distribution of dinucleotides led us to ask whether overall GC content might vary between the two p63RE motif classes, as increasing GC content is linked to increased activity of regulatory elements (Colbran et al., 2017; Lecellier et al., 2018). Unique p63RE have higher overall GC content and these differences are primarily in regions flanking the central p63RE (Fig. 3K). CREs containing p53RE+p63RE motifs are more active despite lower overall GC content relative to unique p63RE. We observe a strong trend between increasing GC content and increasing CRE activity for p53RE+p63RE in the absence of p53 (Fig. 3E,M), which is not observed when p53 is present (Fig. 3D,M). For unique p63 motifs, CRE activity (Fig. 3H,I) closely matches trends in GC content in WT and *TP53* ^-/-^ conditions (Fig. 3L), perhaps suggesting increased GC content between elements can overcome the absence of a strong transactivator like p53.

Taken together, these data suggest that specific classes of p63REs lead to modest differences in p63-dependent CRE transcriptional output. These observations suggest that for elements regulated by p53, motif type and p53 occupancy/affinity are important determinants of high transactivation relative to other sequence-intrinsic features, like GC content. On the contrary, these elements are p63-dependent, but increased p63 enrichment and p63RE motif features do not directly result in higher transactivation.

### Local sequence content and co-occurring transcription factor motifs are associated with differential p63-dependent activation and repression

Our data suggest additional intrinsic DNA sequence characteristics like GC content contribute to maximal activity of p63-bound elements beyond those that directly affect recruitment of p63. p53-bound elements are more active and DNA motifs with higher occupancy lead to higher overall transcriptional output. Our data suggest that loss of p63 binding via mutation of the central p63RE leads to a marked decrease in CRE-driven transcriptional activity. When viewed in aggregate, these results suggest p63 predominantly activates transcription. The ΔNp63ɑ isoform of p63, which is the predominant isoform in basal epithelial cells like MCF-10A, mediates both transcriptional activation and repression, along with chromatin remodeling activities, when binding to regulatory sequences (Bao et al., 2015; Fisher et al., 2020; Yu et al., 2021). We asked whether p63 and p53 transcriptional activation might mask other context dependent activities like repression by examining CRE behavior in the absence of p53. We classified p63-dependent CRE activity as “activating” if mutation of the p63RE led to lower activity relative to WT and “repressing” if lack of p63 binding led to more transcriptional output. Overall, we observe p63-dependent transcriptional activation at nearly 30% (4044/13696) of CREs and repression at only 10% (1345/13696) (Fig. 4A). Interestingly, the activity of most p63-bound elements is not affected when the central p63RE is mutated, regardless of motif type (Fig. 4A). These data potentially suggest that either p63 has non-transcriptional roles that cannot be measured via STARR-seq-style reporter assays or that these elements are strongly cell-type or context-dependent.

**Figure 4.**
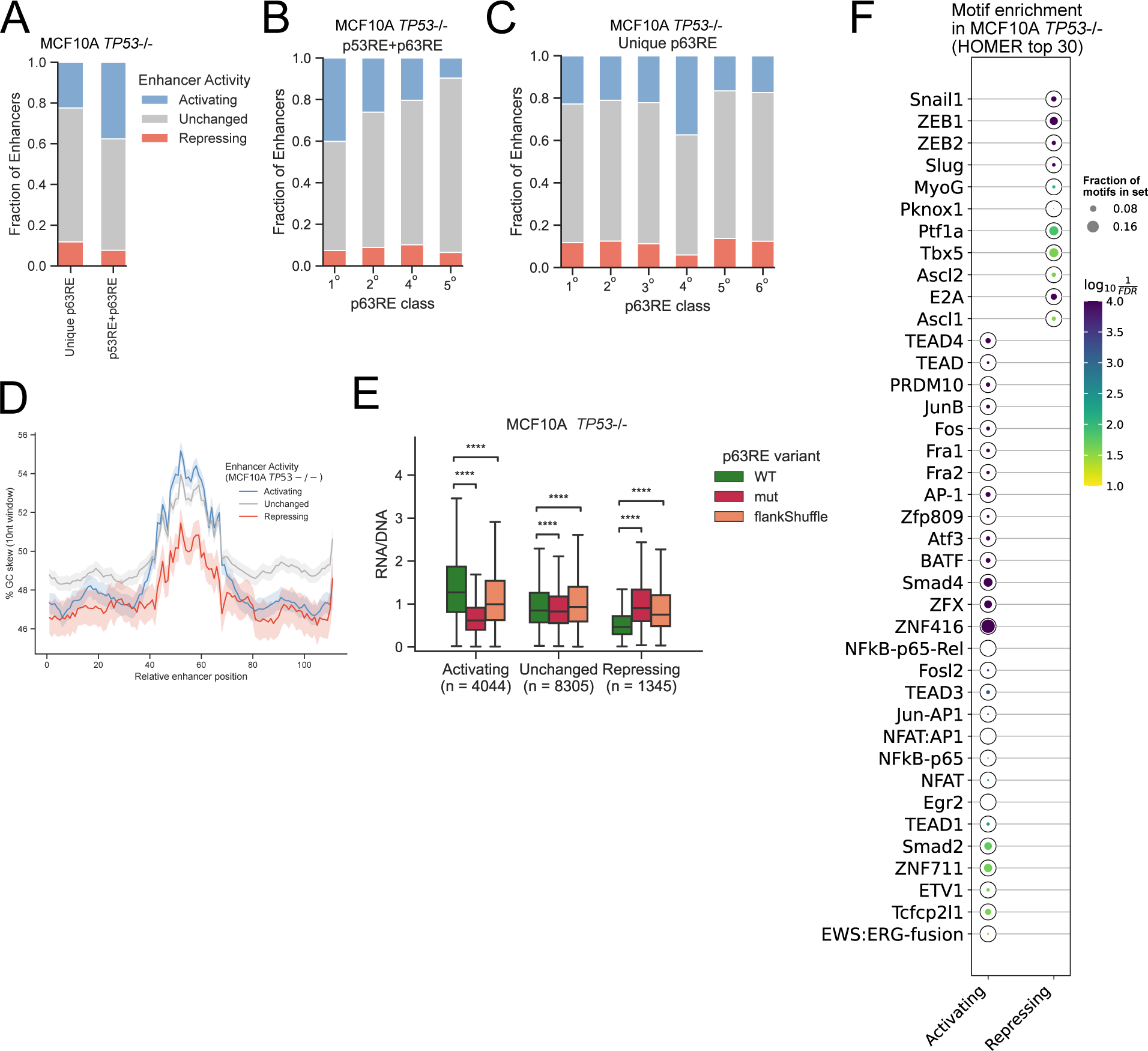
Functional characterization of p63-dependent CRE transcriptional activity. Functional activity is defined by 1.5 fold-change (WT/mut) cutoff where “Activating” is >1.5, “Repressing” is <1.5 and the remaining are defined as “Unchanged”. (A) Distribution of enhancer function in MCF-10A TP53-/- cells in each motif type. Distribution of enhancer function in p53RE+p63RE (B) motif classes or p63 UniqueRE (C) motif classes as defined in Fig. 3A, F. (D) Average GC content across enhancer region by enhancer function. Shaded area represents a 95% confidence interval. (E) WT, mut and flankShuffle enhancer activity in MCF-10A *TP53*-/- cells by function (****: *p*-value < 0.0001, Wilcoxon signed-rank test). (F) Top 30 enriched motifs in “Activating” or “Repressing” enhancer groups relative to “Unchanged” enhancer groups. Motif enrichment was performed using HOMER (Heinz et al., 2010; Duttke et al., 2019). Dot size represents the fraction of CREs containing the specified motif. Color scale indicates Bonferroni-corrected *p*-value.

Activities varied based on the type of p63RE motif found within the CRE. Those containing a p53RE+p63RE motif are almost twice as likely to require p63 for transcriptional activation (Fig. 4A). Primary p53RE+p63RE motifs more frequently lead to p63-dependent activation compared to any other class whereas p63-mediated repression is generally similar across subclasses of p63 response elements (Fig. 4B). Activation at unique p63RE sites is relatively similar across motif types with the exception of those containing quaternary motifs (Fig. 4C). Because they are found in only 65 total regulatory elements, we expect that quaternary motifs are unlikely to broadly represent a general feature that underlies p63-dependent transcriptional activation. GC content spanning the central p63RE is slightly reduced in p63-repressed elements relative to those where p63 activates transcription (Fig. 4D). p63RE type and nucleotide content partially reflect differences between p63-mediated activation and repression, although these effects are relatively modest. Therefore, our data suggest the p63RE motif is critical, but that variation in sequence content between elements is not a major determinant in p63-dependent transcriptional activation and repression.

Cis-regulatory element activity is controlled by the total complement of transcription factors and co-factors interacting with the element (Kulkarni and Arnosti, 2003; Jindal and Farley, 2021). We asked whether sequences, and therefore other DNA binding factors, flanking the central p63RE motif might contribute to activation or repression of p63-bound CREs. As expected, mutation of the central p63RE led to decreased activity at p63-activated elements and an increase in activity at p63-repressed elements, indicative of p63-dependent activity (Fig. 4E). Shuffling sequences flanking the central p63RE led to partial loss of function for both p63-activated and repressed elements. These data suggest that DNA sequences outside of the central p63RE contribute to p63-mediated activities, but the specific mechanisms are not known. We then explored if particular transcription factor motifs might be enriched in CREs where p63 activity was either activating or repressing (Fig. 4F). We used p63RE-containing elements whose activity was p63-independent (unchanged) as the background control to specifically identify unique motifs that might contribute to either activation or repression versus general enrichment with p63RE. p53 family motifs are enriched in activating elements, likely reflecting the observation that most activating elements contain the primary sub-motif which is most closely aligned with the canonical motif models used in the HOMER motif finding algorithm (Table S3) (Heinz et al., 2010; Duttke et al., 2019). Activating elements are also broadly enriched with AP-1 family motifs consistent with these elements supporting transcriptional activation (Biddie et al., 2011; Thurman et al., 2012; Seo et al., 2021). Elements repressed by p63 lack these canonical trans-activator motifs, but are enriched for a series of known transcriptional repressors like Snail, Slug, Zeb1, and Zeb2. All four of these factors have established roles in transcriptional repression during epithelial-to-mesenchymal transition (Peinado et al., 2007; Kalluri and Weinberg, 2009; Pastushenko and Blanpain, 2019). p63-repressed elements are also enriched for motifs for a select set of lineage-specific transcription factors such as Ascl2, MyoG, Tbx5, and Pitf1a, perhaps suggesting p63 and these factors might cooperate to repress key elements during lineage transitions during directed differentiation (Pattison et al., 2018; Li et al., 2019). Although motifs for known activators and repressors are enriched in flanking regions of p63-bound CREs with specific activities, we cannot rule out that DNA shape or p63RE-adjacent context might contribute to changes in p63 binding and activity as they do for p53 (Senitzki et al., 2021; Safieh et al., 2023).

### Cell identity and isoform availability influence p63-bound cis-regulatory element activity

Our data indicate that most p63-bound CREs are not dependent on p63 for their transcriptional activity in MCF-10A STARR-seq assays (Fig. 4A). p63 is a context-dependent transcriptional activator and repressor whose activity is restricted to epithelial cells. While broadly important as a regulator of lineage specification and self-renewal, p63 activity varies across epithelial cell types. For example, p63 is amplified in many squamous cell carcinomas and is associated with poor prognosis and pro-tumorigenic phenotypes (Latil et al., 2017; Saladi et al., 2017; Abraham et al., 2018). We therefore asked whether p63 expression across different epithelial contexts might lead to differential activity of p63-bound CREs in our assay. HaCaT are a spontaneously immortalized keratinocyte line that can undergo squamous differentiation in culture and preserve many of the features of normal human keratinocytes (Wilson, 2013). SCC-25 are squamous cell carcinoma of the tongue, a cancer type that is highly-dependent on p63 for proliferation (Thurfjell et al., 2005; Latil et al., 2017; Saladi et al., 2017; Pokorna et al., 2022). Importantly, both cell lines have inactivating mutations in p53 that limit the analysis to p63-dependent activities (Bamford et al., 2004). HaCaT and SCC-25 also express higher levels of p63 than MCF-10A cells (Fig. 5A) (Sethi et al., 2015), allowing us to ask whether increasing cellular p63 concentration might alter CRE activity.

**Figure 5.**
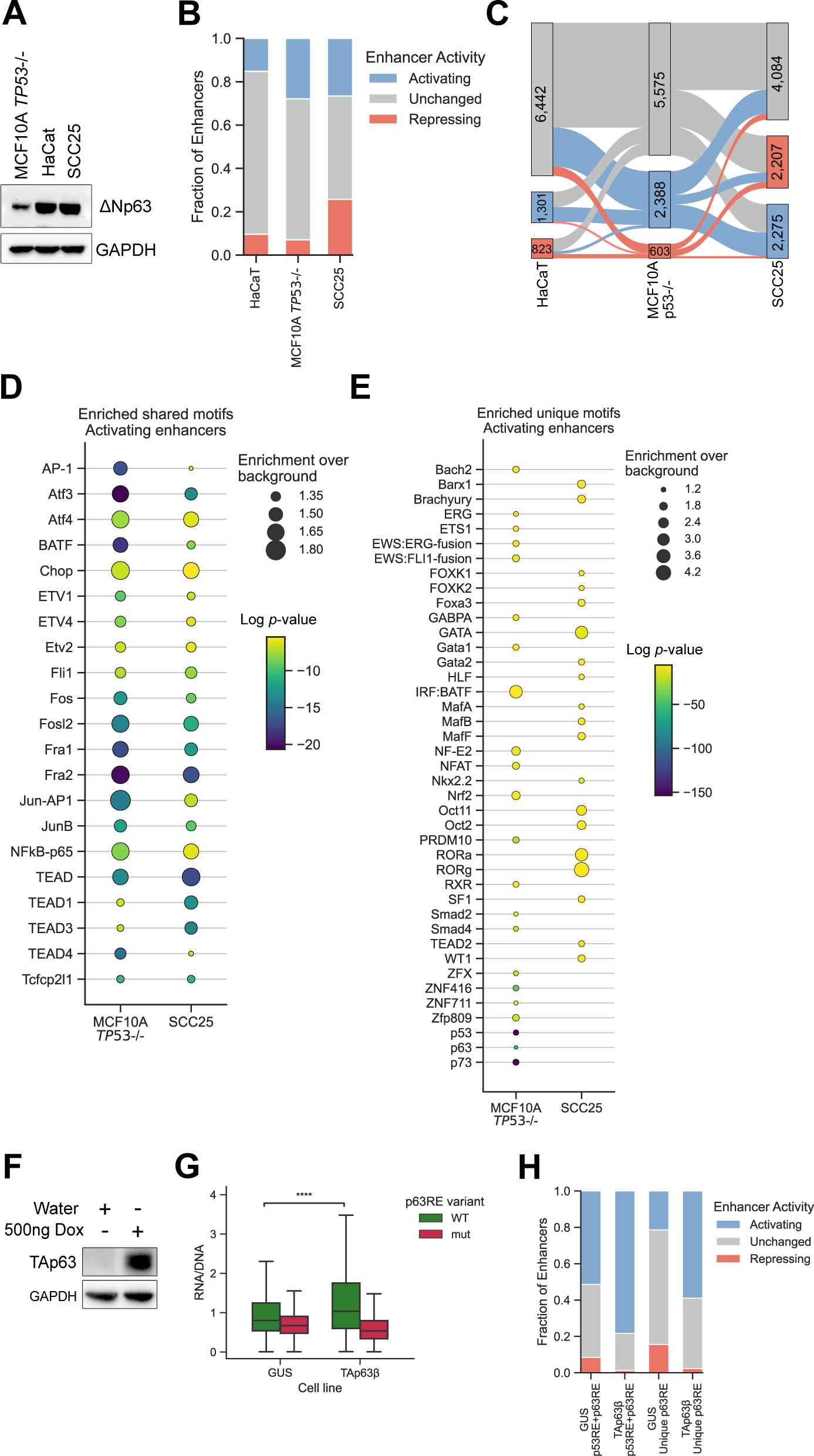
Cell type and context-dependent effect on p63-dependent CRE activity. (A) Western blot analysis of p63 expression in MCF-10A *TP53*-/-, HaCaT and SCC-25 cells. GAPDH is used as a loading control. (B) Distribution of p63-dependent CRE function in MCF-10A *TP53*-/-, HaCat, or SCC-25 cells (N=8,566). (C) Sankey diagram derived from (B) depicting changes in p63-dependent CRE function across three cell types. Numbers indicate CRE counts in each group and cell line. (D-E) Scatter plots highlighting transcription factor motifs enriched in p63-activated CREs that are shared (D) between SCC-25 and MCF-10A *TP53*-/- cell lines or uniquely enriched in each (E). Dot size represents fold-change enrichment over background. Color scale indicates log-transformed *p*-value. (F) Western blot showing Doxycycline-inducible TAp63β or GUS control expression in MCF-10A cells. Cells were treated for 8h with either 500 ng/ml Dox or water as vehicle control. (G) WT and mut CRE activity (N=16,143) in either control (GUS) or TAp63β induced cell line (****: *p*-value < 0.0001, Wilcoxon signed-rank test). (H) Distribution of p63-dependent CRE function in either control or TAp63β induced cell line.

We transfected the MPRA library into either HaCat or SCC-25 cell lines and measured transcriptional output as previously described. Due to differences in transfection efficiency between cell types, we ultimately recovered 8,566 elements with paired WT and mutant expression data across all replicates of MCF-10A *TP53* ^-/-^, HaCaT, and SCC-25 cell lines. We asked whether variation in cell type or p63 expression levels might alter the scope of p63-dependent activation or repression. Overall, the percentage of p63-activated elements is similar, albeit slightly lower, in HaCaT and SCC-25 compared to MCF-10A. SCC-25 cells have nearly 3-fold more p63-repressed elements than the other cell lines (Fig. 5B) although we observe relatively few enriched transcription factor motifs that might contribute to this repression (Table S3). p63-dependent CRE activity varies across these three epithelial cell contexts as most CREs have varied activity across at least two cell lines (Fig. 5C). p63 expression is elevated in both HaCaT and SCC-25, although this does not appear to directly correlate with observed differences in CRE activity. Nearly 30% of CREs do not require the central p63RE for transcriptional activity in any condition (Fig. 5B,C), suggesting they function independent of p63. These results indicate that the collective action of p63 and other transcription factors might underlie the observed variability in gene regulatory element activity.

We then focused on the cell line-specific variability in p63-mediated transcriptional activation. Slightly over 40% (988) of p63-activated CREs are shared between MCF-10A and SCC-25, leaving substantial variability in p63-dependent activity between CREs in SCC-25 (1,287) and MCF-10A (1,400). We therefore examined the differences in transcription factor motif enrichment between p63-dependent CREs in MCF-10A and SCC-25. AP-1 and TEAD family motifs are enriched at p63-activated CREs in both MCF-10A and SCC-25 consistent with their reported roles in transcriptional activation at regulatory elements (Fig. 5D) (Biddie et al., 2011; Currey et al., 2021; Seo et al., 2021). Ets, Smad, and C2H2 zinc finger-related motifs are uniquely enriched in MCF-10A, while SCC-25 p63-activated CREs are enriched for multiple unique transcription factor family motifs, including those from the GATA, FOX, Oct, and Maf families (Fig. 5E). While additional work is needed to determine the extent to which these putative transcription factors are involved, our results provide evidence that p63-dependent transcriptional activity is influenced by the activity of other transcription factors in a cell type-dependent fashion.

Although specific elements shift between being activated or repressed by p63 depending on epithelial cell context, ultimately, p63 is still not required for activity of most p63-bound CREs (Fig. 5C). p53 and p63 share highly overlapping DNA response element motifs (el-Deiry et al., 1992; Perez et al., 2007; Riege et al., 2020), and p53 binding leads to near universal transcriptional activation due to the presence of a strong N-terminal transactivation domain (TAD) (Fischer et al., 2014; Verfaillie et al., 2016). Basal epithelial cells primarily express ΔNp63ɑ (Fig. 5A), but multiple N- and C-terminal isoforms of p63 can be expressed in different cellular contexts (Marshall et al., 2021). We therefore asked whether the p63-bound CRE activity might have p63 isoform-specific dependence. TAp63 isoforms are expressed in late stages of keratinocyte differentiation and are required for the response to genotoxic damage in germ cells (Koster et al., 2004; Truong et al., 2006; Beyer et al., 2011; Deutsch et al., 2011). TAp63ɑ drives high levels of transcriptional activation in a stimulus-dependent fashion (Deutsch et al., 2011; Coutandin et al., 2016), whereas other C-terminal isoforms, like TAp63β are constitutively active (Lena et al., 2021). We therefore chose to measure CRE activity in response to expression of TAp63β in order to avoid potential crosstalk with genotoxic or other cell stress pathways. We transfected the MPRA library into MCF-10A cells where either a control protein (GUS) or TAp63β was expressed under doxycycline-inducible control (Fig. 5F). The distribution of p63-bound CREs in control conditions that are p63-activated, repressed, or independent is highly similar to our previous assays in MCF-10A and MCF-10A *TP53* ^-/-^ cell lines (Fig. 5H vs. Fig. 4A). TAp63β expression led to increased overall CRE activity and importantly, this activity is dependent on the central p63RE motif (Fig. 5G). Nearly 70% of CREs display p63RE-dependent transcriptional activation when TAp63β is expressed, more than a 2-fold increase compared to control conditions (Fig. 5H). This is most observable at CREs with unique p63RE which see a 3 fold shift towards p63-dependent transcriptional activation and a near-complete loss of repression in the presence of TAp63β. Taken together, these results suggest that context-dependent p63 isoform expression alters the activity of cis-regulatory elements and that TA-isoforms primarily activate transcription.

## Discussion

The importance of p63 in regulating epithelial cell identity is supported by extensive genetic and biochemical evidence. p63 regulates epidermal development through its transcription factor activity and control of epithelial-specific transcription and chromatin structure. These activities require p63 binding and context-specific transcriptional regulation, but how DNA sequence at regulatory elements affects p63 activity is an open question. Here, we examine whether the sequence content and context of p63 binding sites controls p63-dependent transcriptional activity using massively parallel reporter assays. We find that sequence content of p63 response element motifs influences p63 binding and transcriptional activity, but that this relationship is complicated. The complex relationship between sequence and function is partially due to p63 roles as both a context-dependent transcriptional activator and repressor.

ΔNp63-dependent repression is relatively rare compared to activation, as has been suggested by studies combining p63 binding and global transcriptome analyses (Riege et al., 2020). Repression can be mediated by C-terminal recruitment of known co-repressors like histone deacetylases or through antagonism of other transcription factors, like p53 (LeBoeuf et al., 2010; Ramsey et al., 2011; Woodstock et al., 2021). Our data suggest that slight variation in p63RE motif sequence content, like varying GC and dinucleotide content, might partially explain varying activation or repression, but this is likely only a minor contributor. Motifs for known repressive transcription factors like Snail, Slug, Zeb1, and Zeb2 are specifically enriched in p63-repressed elements. Interestingly, these factors are key regulators of epithelial-to-mesenchymal transition (EMT) (Peinado et al., 2007), a process globally suppressed by p63 (Yoh et al., 2016; Latil et al., 2017). While they may antagonize each other globally, p63 and these EMT-promoting factors may have cooperative roles in repression of specific genes. p63 switches between repressive and activating states during development, starting by repressing non-epithelial lineage genes before switching to activation during epithelial commitment (Pattison et al., 2018; Santos-Pereira et al., 2019). p63 also locally represses some TFAP2C binding sites important for early epidermal specification during later stages of keratinocyte maturation (Li et al., 2019). Repression in these settings results from p63-dependent alteration of local chromatin structure or chromatin modification by HDACs which may not be directly measured using plasmid-based MPRA style assays. The full scope of p63-mediated repression, and the extent of its regulation by DNA sequence alone, might not be observable in a single terminally-differentiated cell line using only MPRA tools.

The relationship between sequence identity and transcriptional output for p63-bound elements is also complicated by context-dependent activity of other transcription factors binding the same p63RE motif. The strongest predictor of regulatory element-driven transcriptional output from our results is the presence of p63RE motifs capable of binding the p63 paralog p53 (Fig. 2D). These elements are highly active and dependent on the central p53/p63 response element, which was recently identified as the strongest predictor of regulatory element-driven transcription (Sahu et al., 2022). The relative activity was preserved in the absence of p53 suggesting that p63 can also drive high-level transcriptional activation (Fig. 2I)(Fig. 4A). How, though, these motifs drive higher expression by p63 is still unclear. Motif identity is linked to higher enrichment and transcriptional output by p53 (Fig. 3C,D), but we did not observe any such relationship for p63. Sites with higher p63 enrichment are not neccessarily more active, as has been observed directly for p53 (Trauernicht et al., 2023). Rather, our data suggest that ubiquity of p63 binding across cell types better reflects increased transcriptional activity (Fig. 2B). Other features, like increasing local H3K27ac (Kouwenhoven et al., 2015; Qu et al., 2018) and transcription factor motifs flanking p63REs, including those for traditional transcriptional activators, contribute to p63-dependent trans-activation (Yang et al., 2006; McDade et al., 2012; Sethi et al., 2017). Craniofacial development requires specific and combinatorial activity of p63 and other transcription factors at an enhancer for *IRF6* (Rahimov et al., 2008; Fakhouri et al., 2012, 2014). Our results on p63RE affinity and occupancy are consistent with a model that enhancers often contain suboptimal binding sites and use motif grammar and syntax to drive appropriate developmental and stimulus-dependent behaviors (Crocker et al., 2015; Farley et al., 2015, 2016; Lim et al., 2024).

p53 is generally regarded as a universal activator of transcription, whereas p63 either activates or represses in a context-dependent manner (Fig. 6A) (Fischer et al., 2014). Our data provide insight into how sequence context, including various sub-classes of the core p63RE motif and flanking transcription factor motifs, can affect these p63-dependent functions. The mechanisms controlling this switch between activities, including when p63 serves as a pioneer or bookmarking factor (Bao et al., 2015; Kouwenhoven et al., 2015), are not fully understood. p63 is also a *bona fide* pioneer transcription factor and controls accessibility at epithelial-specific regulatory elements (Kouwenhoven et al., 2015; Pattison et al., 2018; Qu et al., 2018; Karsli Uzunbas et al., 2019; Li et al., 2019; Lin-Shiao et al., 2019; Santos-Pereira et al., 2019; Yu et al., 2021). Massively parallel reporter assays are powerful tools to study transcriptional activation and repression, but their design can limit the range of transcription factor activities that can be directly measured (Inoue and Ahituv, 2015; Trauernicht et al., 2020). Most elements display p63 expression-dependent chromatin accessibility in epithelial cell types (Fig. 1F) but do not rely on ΔNp63ɑ for their observed transcriptional activity (Fig. 4A). Roles for p63 in enhancer:promoter interactions, such as those observed during p63-dependent directed keratinocyte differentiation (Pattison et al., 2018; Li et al., 2019; Qu et al., 2019), would be difficult to measure in a non-genomic context. One other possibility to be investigated in future studies is that many elements require p63 for *in vivo* chromatin accessibility but not for direct transcriptional activation or repression. Complementary approaches, like genome-scale MPRA and loci-specific genetic dissection, are likely required to fully unravel the range of p63-dependent activities at regulatory elements.

**Figure 6.**
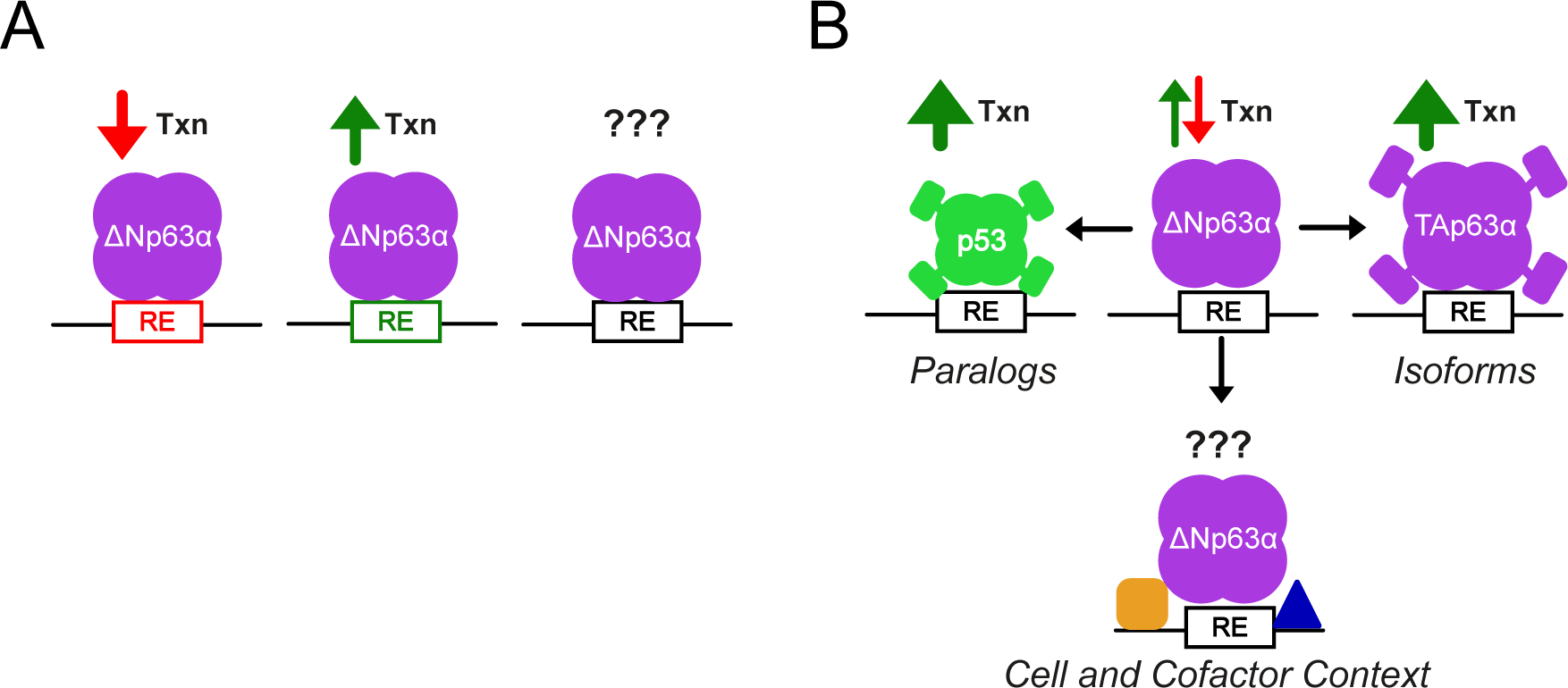
(A) Schematic illustrating differential activity of ΔNp63ɑ at *cis*-regulatory elements. While sequences can elicit either ΔNp63ɑ-dependent transcriptional activation or repression, many appear ΔNp63ɑ-independent. (B) Schematic depicting how p63 paralogs and isoforms binding to the same element can elicit different outcomes compared to ΔNp63ɑ, including high level transcriptional activation by p53 and TAp63. Other p63 paralogs and isoforms may have different activities depending on the presence of particular protein domains, and are not depicted here. ΔNp63ɑ activity at *cis*-regulatory elements also can change based on cell context or on the availability of local transcription factors, although the molecular grammar of these interactions and how they give rise to differential function remain unclear.

The seeming lack of p63-dependent transcriptional control at a substantial number of p63-bound regulatory elements led us to ask whether cell context might drive differential p63 activities. Enhancers are well-known to exhibit cell type and context-dependent activities controlled by variable expression of transcription factors and co-factors (Spitz and Furlong, 2012; Heinz et al., 2015). p63 expression is strongly lineage restricted during development and homeostasis and varied p63 levels have been linked to human cancers (Massion et al., 2003; Graziano and De Laurenzi, 2011; Tucci et al., 2012; Pickering et al., 2014; Saladi et al., 2017). Elevated p63 expression is strongly linked to pro-survival pathways in squamous cell carcinomas (SCC) (Thurfjell et al., 2005; Ramsey et al., 2013; Abraham et al., 2018). These collective observations led us to investigate whether p63-dependent regulatory element behavior varied across epithelial cell contexts. SCC-25, a head and neck squamous cell carcinoma cell line, in particular showed varied activity of p63-dependent regulatory elements (Fig. 5B,C). Although more elements displayed p63-dependent repression than in MCF-10A, these elements lacked specific enrichment for transcription factor motifs that might cooperate with p63 to reduce transcriptional output (Table S3). In contrast, p63-dependent activation in SCC-25 was coupled with enrichment of different TF motifs than those associated with activation in MCF10A including a range of factors with known epithelial functions, like the Maf and Forkhead families (Fig. 5E) (Lopez-Pajares et al., 2015; Napoli et al., 2024). Regulatory elements with cell-specific activities appear to utilize different combinations of co-enriching motifs alongside p63 (Donohue et al., 2022). The extent to which p63 amplification or other transcription factor availability drives differential p63-dependent activities at gene regulatory elements, both during development and disease, requires more investigation to unravel. This might include combining advances in genome-scale reporter assays, single-cell spatial transcriptomics, and machine-learning assisted design that have led to the ability to design synthetic enhancers with defined cell type-specific activities (Taskiran et al., 2024).

Transcription factors are commonly spliced to produce various isoforms which often display differential activities (Wang et al., 2008; Lambourne et al., 2024). The role of p63 at regulatory elements is further complicated by the complexities of transcription factor isoforms and paralogs. The constitutively active TAp63β isoform activated transcription at most regulatory elements (Fig. 5H) similar to near-universal activation by p53 (Verfaillie et al., 2016; Peng et al., 2020; Sahu et al., 2022). TAp63ɑ is critical in the germline and in late keratinocyte differentiation and TAp63ɑ has stimulus-dependent activity unlike ΔNp63 isoforms (Koster et al., 2004; Truong et al., 2006; Beyer et al., 2011; Deutsch et al., 2011). Gene regulatory elements activated uniquely by TAp63 isoforms are bound by both TA and ΔNp63 isoforms. The role of ΔN isoforms at these elements then may be to establish and bookmark local chromatin structure for later TA isoform activity, as ΔNp63 can do for regulated cell lineage-specific p53 activity (Karsli Uzunbas et al., 2019). The interplay between transcription factor paralogs with overlapping DNA binding activity can drive variable gene regulatory element activity (Fig. 6B). p53 and p73 share considerable tissue expression and binding site overlap with p63 (Marshall et al., 2021), so the extent to which paralog expression influences p63-dependent behaviors should not be overlooked. Similarly, p73 may also influence p63 activity through formation of mixed p63:p73 heterotetramers suggesting yet another mechanism influencing regulatory element behavior (Strubel et al., 2023). Our results suggest that significant additional effort should be placed into identifying how cell type, developmental stage, or stimulus-dependent conditions might lead to p63 isoform switching, paralog expression, and, ultimately, varied p63-dependent gene regulatory activity (Fig. 6B).

In conclusion, we present a near genome-scale analysis of p63-dependent regulatory element activity. Our data are consistent with varying roles of p63 in literature and suggest that while sequence content is important, other local cofactors, isoform switching, paralog expression, and chromatin are critical context-dependent regulators of p63-dependent CRE activity. Unraveling the full scope of p63 activities will likely require multiple complementary approaches including specific assays focused on p63-dependent chromatin remodeling, native approaches for examining sequence content such as genome editing, and new computational tools like AI and deep learning.

## MATERIALS AND METHODS

### Cell culture

All human mammary epithelial cell lines MCF-10A TP53+/+ and TP53-/- (Sigma-Millipore clls1049) were cultured in 1:1 Dulbecco’s Modified Eagle’s Medium: Ham’s F-12 (Gibco, #11330-032), supplemented with 5% Horse Serum, (Gibco, #16050-122), 20 ng/mL epidermal growth factor (Peprotech, #AF-100-15), 0.5 µg/mL hydrocortisone (Sigma, #H-0888), 100 ng/mL cholera toxin (Sigma, #C-8052), 10 µg/mL insulin (Sigma, #I-1882), and 1% penicillin-streptomycin (Gibco, #15240-062). MCF10A TP53-/- cells were obtained from Sigma-Millipore (clls1049) and were cultured Human HNSCC cell line SCC-25 (kind gift of C. Michael DiPersio, Albany Medical College) were cultured in 1:1 Dulbecco’s Modified Eagle’s Medium: Ham’s F-12, supplemented with 10% FBS (Corning, #35-016-CV), 1% penicillin-streptomycin and 400 ng/ml hydrocortisone. Human transformed keratinocyte cell line HaCat and HEK293FT cells were cultured in Dulbecco’s Modified Eagle’s Medium 1X (Corning 10-013-CV) and supplemented with 10% FBS and 1% penicillin-streptomycin. All cell lines were cultured at 37°C and 5% CO_2_.

### Lentiviral Production

Lentiviral particles were packaged by transfecting 600ng psPAX2, 400ng of pMD2.G, and 1ug of pCW57.1 containing either TAp63β or β-glucoronidase (GUS) control in HEK293FT cells at a density of 600,000 per well. pCW57.1 (pCW57.1 was a gift from David Root, Addgene plasmid # 41393 ; http://n2t.net/addgene:41393 ; RRID:Addgene_41393), psPAX2, and pMD2.G (psPAX2 and pMD2.G were a gift from Didier Trono, Addgene plasmid # 12260 ; http://n2t.net/addgene:12260 ; RRID:Addgene_12260) were obtained from Addgene. GUS control plasmid was provided as part of the LR Clonase II enzyme kit (Invitrogen 11791020). Lentiviral supernatants were collected at 24 and 48 hours and concentrated via spin dialysis. Viral supernatants were added to MCF-10A cells with 8ug/ml polybrene for 24 hours and then replaced with fresh media. 2ug/ml puromycin was added 48 hours after infection and cells were selected for 72 hours.

### Western blotting

Protein was isolated using custom made RIPA buffer (50 mM Tris-HCl pH 7.4, 150 mM NaCl, 1% NP-40, 0.5% sodium deoxycholate, 0.1% SDS, 1mM EDTA, 1% Triton x-100) supplemented with protease inhibitor (Pierce, 78442). Concentration of isolated protein was measured using a microBCA kit (Pierce, 23227) and 25µg was loaded on a 4-12% Bis-Tris protein gel (Invitrogen, NP0321BOX). Protein size was analyzed using PageRuler™ Prestained Protein Ladder (Thermo 26616). Membranes were blocked in 5% non-fat milk in TBS-T. Antibodies used included rabbit anti-ΔNp63 antibody (Cell Signaling E6Q3O), mouse anti-p53 (BD Biosciences 554293), mouse anti-TAp63 (BioLegend 938102) rabbit anti-GAPDH antibody (Cell Signaling D16H11), and rabbit anti-!-Glucuronidase (Sigma Aldrich G5545).

### Massively parallel reporter assay (MPRA) design

MPRA query regions were selected from a recent analysis of multiple p63 ChIP-seq datasets (Riege et al., 2020). Only p63 binding events observed in 8 or more independent experiments and containing p63 response element (p63RE) sequences (17,310 locations) were considered for analysis due to DNA synthesis constraints. MPRA regions were centered on the p63RE and were extended to a total length of 119 or 120 bp based on the length of the p63RE. Genomic coordinates corresponding to each p63 MPRA region were used to extract DNA sequence information from the hg38 UCSC genome assembly using bedtools. Either the entire MPRA sequence (full shuffle), the p63RE (shuffle), or the regions flanking the p63RE (flank shuffle) were randomly scrambled, while preserving GC content, to produce three variants. Position weight matrices for the p63RE were generated using consensusMatrix (Biostrings R package) and visualized using seqLogo. A fourth variant (mutant) was designed where consensus nucleotides found within the p63RE at a frequency greater than 75% were substituted to preserve GC content. A schematic of all substitutions can be found in Figure 1A. Adapter sequences were then added to the 5’ (5’-TCCCTACACGACGCTCTTCCGATCT) and 3’ (5’-AGATCGGAAGAGCACACGTCTGAAC) end of each MPRA sequence. Sequences for all MPRA regions can be found in Table S1. An oligo pool containing all 86,550 reporter sequences was synthesized by GenScript (Piscataway, NJ, USA).

### Cloning oligo pool library

MPRA oligo plasmid pool was cloned as described with the following adjustments (Neumayr et al., 2022). Plasmid backbone pGB118, with added Illumina i5/i7 sequences flanking the cloning site, was digested with *AgeI* and *SalI* as described (Baniulyte et al., 2023). pGB118 was based on hSTARR-seq_ORI vector, which was a gift from Alexander Stark (Addgene plasmid # 99296 ; http://n2t.net/addgene:99296 ; RRID:Addgene_99296). The oligo pool was amplified using Q5 polymerase and SL1947 (5’-TCCCTACACGACGCTCTTC) and SL1948 (5’-GTTCAGACGTGTGCTCTTC) primers for 15 cycles. Amplicons were then cloned into pGB118 using HiFi assembly (NEB M0492S) and HiFi reactions were transformed into DH5alpha cells (NEB C2987H) and grown in LB culture. Plasmid library was purified using ZymoPURE Gigaprep kit (Zymo D4204).

### Plasmid pool transfection, library preparation and sequencing

Two biological replicates were performed for each cell type and condition at approximately 50 million cells per replicate and one biological replicate was performed for isoform (GUS, TAp63β) overexpression assay. 10 μg of plasmid library were transfected per 5 million cells via lipofection (Polyplus #101000046, #101000025). For TAp63β inducible cell line and negative control (GUS) cell line, doxycycline was added at 500 ng/ml at the same time as transfection. Cells were harvested after 24 hours and total RNA was extracted (Quick RNA, Zymo, #R1055). 30 μg per replicate of poly-adenylated mRNA was isolated using oligo d(T) beads (NEB E7490L). Resulting mRNA was split into 6 separate reactions and cDNA was synthesized using a gene-specific primer (5’-CTCATCAATGTATCTTATCATGTCTG-3’) and MultiScribe Reverse Transcriptase (Invitrogen, 4311235). Following cDNA synthesis, all cDNA samples were pooled and one Junction-PCR reaction was performed with 16 cycles (5’-TCGTGAGGCACTGGGCAGGTGTC,CTTATCATGTCTGCTCGAAGC-3’) and i5 and i7 primers from NEBNext Oligo Kit (E7600S) were used for Illumina barcoding with between 5 and 9 cycles. MPRA plasmid DNA pools were amplified with i5 and i7 primers as a control for oligo representation. All libraries were pooled and sequenced as single-end, 100bp on an Illumina NextSeq 2000 instrument at the University at Albany Center for Functional Genomics.

### MPRA library data analysis

FASTQ files for plasmid DNA and enhancer RNA libraries were mapped using exact pattern match using custom Python scripts. Enhancers that had less than 5 reads were removed. Raw read count table is available under Gene Expression Omnibus accession number GSE266670. Additionally, for Figures 1-4 were MCF-10A and MCF-10A *TP53*-/- cell lines were used, only enhancers that had a match for every enhancer variant (WT, mut, shuffle, flankShuffle, fullShuffle) were kept (n = 13,696). Where MCF-10A *TP53*-/-, HaCaT and SCC-25 (Fig. 5 B-E) or MCF-10A TAp63β and GUS overexpression cell lines were considered, enhancers that only had a WT and mut matched variants were kept (N = 8,566 and N = 16,143, respectively). All reads were normalized to the total number of reads per sample and expression values are represented as RNA/DNA ratio and averaged between the replicates. Normalized expression values and enhancers considered for each figure are listed in Table S2. Statistical tests were performed using Python packages (SciPy, scikit-posthocs).

### Data Integration

MCF-10A p63, p53, H3K27ac, H3K4me2, and H3K4me3 datasets were downloaded from Gene Expression Omnibus accession GSE111009 (Karsli Uzunbas et al., 2019). Raw data were mapped to the GRCh38 reference assembly using hisat2 and biological replicates were combined using the merge function of samtools (Li et al., 2009; Kim et al., 2019). Enrichment at MPRA regions was quantified and visualized using deepTools (Ramírez et al., 2016). p53 and p63 enrichment data were quantified within 1bp bins across the entire p53/p63RE motif location and heatmaps were generated using 10bp bins across a region spanning -/+ 1,000 bp from the p53/p63RE motif. Datasets used to integrate ENCODE ChromHMM, candidate Cis Regulatory Elements (cCRE), and DNase Hypersensitivity Clusters (DHS) with MPRA genomic locations were obtained from repositories as specified in Table S4. Intersections between p63 CRE regions and various datasets were performed using either bedTools or BigBedtoBed (Kent et al., 2010; Quinlan and Hall, 2010). Motif enrichment analyses were performed using HOMER (Heinz et al., 2010; Duttke et al., 2019). GC and dinucleotide content analyses were performed using custom nucleotide counting scripts in Python.

### Data availability

MPRA datasets from this manuscript are available under Gene Expression Omnibus (GEO) accession GSE266670.

## Acknowledgments

This work was supported by NIH R35 GM138120 to MAS. DLW was partially supported by NIH T32 GM132066. The authors would like to thank the University at Albany Center for Functional Genomics for sequencing support and the RNA Institute at the University at Albany for additional equipment support and for generous support of the trainees on this manuscript.

---

**Table S1.** Relevant CRE information and sequences used in this study.

**Table S2.** Normalized expression values (RNA/DNA) for cell type- and CRE variant-matched enhancers used in this study.

**Table S3.** HOMER knownResults output for “Activating” or “Repressing” CREs in MCF-10A *TP53*-/- and SCC-25 cell lines.

**Table S4.** Data sources.

## Bibliography

Abraham, C. G., Ludwig, M. P., Andrysik, Z., Pandey, A., Joshi, M., Galbraith, M. D., et al. (2018). ΔNp63α Suppresses TGFB2 Expression and RHOA Activity to Drive Cell Proliferation in Squamous Cell Carcinomas. Cell Reports 24, 3224–3236. doi: 10.1016/j.celrep.2018.08.058

Amiel, J., Bougeard, G., Francannet, C., Raclin, V., Munnich, A., Lyonnet, S., et al. (2001). TP63 gene mutation in ADULT syndrome. Eur J Hum Genet 9, 642–645. doi: 10.1038/sj.ejhg.5200676

Arnold, C. D., Gerlach, D., Stelzer, C., Boryn, L. M., Rath, M., and Stark, A. (2013). Genome-wide quantitative enhancer activity maps identified by STARR-seq. Science 339, 1074–7. doi: 10.1126/science.1232542

Bamford, S., Dawson, E., Forbes, S., Clements, J., Pettett, R., Dogan, A., et al. (2004). The COSMIC (Catalogue of Somatic Mutations in Cancer) database and website. Br J Cancer 91, 355–358. doi: 10.1038/sj.bjc.6601894

Baniulyte, G., Durham, S. A., Merchant, L. E., and Sammons, M. A. (2023). Shared Gene Targets of the ATF4 and p53 Transcriptional Networks. Molecular and Cellular Biology 0, 1–24. doi: 10.1080/10985549.2023.2229225

Bao, X., Rubin, A. J., Qu, K., Zhang, J., Giresi, P. G., Chang, H. Y., et al. (2015). A novel ATAC-seq approach reveals lineage-specific reinforcement of the open chromatin landscape via cooperation between BAF and p63. Genome Biology 16, 284. doi: 10.1186/s13059-015-0840-9

Barral, A., and Zaret, K. S. (2023). Pioneer factors: roles and their regulation in development. Trends in Genetics 0. doi: 10.1016/j.tig.2023.10.007

Beyer, U., Moll-Rocek, J., Moll, U. M., and Dobbelstein, M. (2011). Endogenous retrovirus drives hitherto unknown proapoptotic p63 isoforms in the male germ line of humans and great apes. Proc Natl Acad Sci U S A 108, 3624–3629. doi: 10.1073/pnas.1016201108

Biddie, S. C., John, S., Sabo, P. J., Thurman, R. E., Johnson, T. A., Schiltz, R. L., et al. (2011). Transcription factor AP1 potentiates chromatin accessibility and glucocorticoid receptor binding. Mol. Cell 43, 145–155. doi: 10.1016/j.molcel.2011.06.016

Bougeard, G., Hadj-Rabia, S., Faivre, L., Sarafan-Vasseur, N., and Frébourg, T. (2003). The Rapp–Hodgkin syndrome results from mutations of the TP63 gene. Eur J Hum Genet 11, 700–704. doi: 10.1038/sj.ejhg.5201004

Candi, E., Rufini, A., Terrinoni, A., Dinsdale, D., Ranalli, M., Paradisi, A., et al. (2006). Differential roles of p63 isoforms in epidermal development: selective genetic complementation in p63 null mice. Cell Death Differ 13, 1037–1047. doi: 10.1038/sj.cdd.4401926

Castro-Mondragon, J. A., Riudavets-Puig, R., Rauluseviciute, I., Berhanu Lemma, R., Turchi, L., Blanc-Mathieu, R., et al. (2022). JASPAR 2022: the 9th release of the open-access database of transcription factor binding profiles. Nucleic Acids Research 50, D165– D173. doi: 10.1093/nar/gkab1113

Celli, J., Duijf, P., Hamel, B. C. J., Bamshad, M., Kramer, B., Smits, A. P. T., et al. (1999). Heterozygous Germline Mutations in the p53 Homolog p63 Are the Cause of EEC Syndrome. Cell 99, 143–153. doi: 10.1016/S0092-8674(00)81646-3

Colbran, L. L., Chen, L., and Capra, J. A. (2017). Short DNA sequence patterns accurately identify broadly active human enhancers. BMC Genomics 18, 536. doi: 10.1186/s12864-017-3934-9

Coutandin, D., Osterburg, C., Srivastav, R. K., Sumyk, M., Kehrloesser, S., Gebel, J., et al. (2016). Quality control in oocytes by p63 is based on a spring-loaded activation mechanism on the molecular and cellular level. eLife 5, e13909. doi: 10.7554/eLife.13909

Crocker, J., Abe, N., Rinaldi, L., McGregor, A. P., Frankel, N., Wang, S., et al. (2015). Low affinity binding site clusters confer hox specificity and regulatory robustness. Cell 160, 191–203. doi: 10.1016/j.cell.2014.11.041

Currey, L., Thor, S., and Piper, M. (2021). TEAD family transcription factors in development and disease. Development 148, dev196675. doi: 10.1242/dev.196675

Deutsch, G. B., Zielonka, E. M., Coutandin, D., Weber, T. A., Schäfer, B., Hannewald, J., et al. (2011). DNA Damage in Oocytes Induces a Switch of the Quality Control Factor TAp63α from Dimer to Tetramer. Cell 144, 566–576. doi: 10.1016/j.cell.2011.01.013

Donohue, L. K. H., Guo, M. G., Zhao, Y., Jung, N., Bussat, R. T., Kim, D. S., et al. (2022). A cis-regulatory lexicon of DNA motif combinations mediating cell-type-specific gene regulation. Cell Genom 2, 100191. doi: 10.1016/j.xgen.2022.100191

Duttke, S. H., Chang, M. W., Heinz, S., and Benner, C. (2019). Identification and dynamic quantification of regulatory elements using total RNA. Genome Res. doi: 10.1101/gr.253492.119

el-Deiry, W. S., Kern, S. E., Pietenpol, J. A., Kinzler, K. W., and Vogelstein, B. (1992). Definition of a consensus binding site for p53. Nat Genet 1, 45–49. doi: 10.1038/ng0492-45

Ernst, J., and Kellis, M. (2015). Large-scale imputation of epigenomic datasets for systematic annotation of diverse human tissues. Nat. Biotechnol. 33, 364–376. doi: 10.1038/nbt.3157

Fakhouri, W. D., Rahimov, F., Attanasio, C., Kouwenhoven, E. N., Ferreira De Lima, R. L., Felix, T. M., et al. (2014). An etiologic regulatory mutation in IRF6 with loss- and gain-of-function effects. Hum Mol Genet 23, 2711–20. doi: 10.1093/hmg/ddt664

Fakhouri, W. D., Rhea, L., Du, T., Sweezer, E., Morrison, H., Fitzpatrick, D., et al. (2012). MCS9.7 enhancer activity is highly, but not completely, associated with expression of Irf6 and p63. Dev Dyn 241, 340–349. doi: 10.1002/dvdy.22786

Farley, E. K., Olson, K. M., Zhang, W., Brandt, A. J., Rokhsar, D. S., and Levine, M. S. (2015). Suboptimization of developmental enhancers. Science 350, 325–328. doi: 10.1126/science.aac6948

Farley, E. K., Olson, K. M., Zhang, W., Rokhsar, D. S., and Levine, M. S. (2016). Syntax compensates for poor binding sites to encode tissue specificity of developmental enhancers. Proc Natl Acad Sci U S A 113, 6508–6513. doi: 10.1073/pnas.1605085113

Fischer, M., Steiner, L., and Engeland, K. (2014). The transcription factor p53: not a repressor, solely an activator. Cell Cycle 13, 3037–3058. doi: 10.4161/15384101.2014.949083

Fisher, M. L., Balinth, S., and Mills, A. A. (2020). p63-related signaling at a glance. J Cell Sci 133. doi: 10.1242/jcs.228015

Fletcher, R. B., Prasol, M. S., Estrada, J., Baudhuin, A., Vranizan, K., Choi, Y. G., et al. (2011). p63 Regulates Olfactory Stem Cell Self-Renewal and Differentiation. Neuron 72, 748–759. doi: 10.1016/j.neuron.2011.09.009

Gebel, J., Tuppi, M., Krauskopf, K., Coutandin, D., Pitzius, S., Kehrloesser, S., et al. (2017). Control mechanisms in germ cells mediated by p53 family proteins. Journal of Cell Science 130, 2663–2671. doi: 10.1242/jcs.204859

Graziano, V., and De Laurenzi, V. (2011). Role of p63 in cancer development. Biochimica et Biophysica Acta (BBA) - Reviews on Cancer 1816, 57–66. doi: 10.1016/j.bbcan.2011.04.002

Hager, G. L., McNally, J. G., and Misteli, T. (2009). Transcription Dynamics. Molecular Cell 35, 741–753. doi: 10.1016/j.molcel.2009.09.005

Halfon, M. S. (2020). Silencers, Enhancers, and the Multifunctional Regulatory Genome. Trends in Genetics 36, 149–151. doi: 10.1016/j.tig.2019.12.005

Heinz, S., Benner, C., Spann, N., Bertolino, E., Lin, Y. C., Laslo, P., et al. (2010). Simple combinations of lineage-determining transcription factors prime cis-regulatory elements required for macrophage and B cell identities. Mol. Cell 38, 576–589. doi: 10.1016/j.molcel.2010.05.004

Heinz, S., Romanoski, C. E., Benner, C., and Glass, C. K. (2015). The selection and function of cell type-specific enhancers. Nature Reviews Molecular Cell Biology 16, 144–154. doi: 10.1038/nrm3949

Inoue, F., and Ahituv, N. (2015). Decoding enhancers using massively parallel reporter assays. Genomics 106, 159–164. doi: 10.1016/j.ygeno.2015.06.005

Janky, R., Verfaillie, A., Imrichová, H., Van de Sande, B., Standaert, L., Christiaens, V., et al. (2014). iRegulon: From a Gene List to a Gene Regulatory Network Using Large Motif and Track Collections. PLoS Computational Biology 10, e1003731. doi: 10.1371/journal.pcbi.1003731

Jindal, G. A., and Farley, E. K. (2021). Enhancer grammar in development, evolution, and disease: dependencies and interplay. Developmental Cell 56, 575–587. doi: 10.1016/j.devcel.2021.02.016

Kalluri, R., and Weinberg, R. A. (2009). The basics of epithelial-mesenchymal transition. J Clin Invest 119, 1420–1428. doi: 10.1172/JCI39104

Karsli Uzunbas, G., Ahmed, F., and Sammons, M.A. (2019). Control of p53-dependent transcription and enhancer activity by the p53 family member p63. J. Biol. Chem. doi: 10.1074/jbc.RA119.007965

Katoh, I., Maehata, Y., Moriishi, K., Hata, R.-I., and Kurata, S. (2019). C-terminal α Domain of p63 Binds to p300 to Coactivate β-Catenin. Neoplasia 21, 494–503. doi: 10.1016/j.neo.2019.03.010

Kent, W. J., Zweig, A. S., Barber, G., Hinrichs, A. S., and Karolchik, D. (2010). BigWig and BigBed: enabling browsing of large distributed datasets. Bioinformatics 26, 2204–2207. doi: 10.1093/bioinformatics/btq351

Kim, D., Paggi, J. M., Park, C., Bennett, C., and Salzberg, S. L. (2019). Graph-based genome alignment and genotyping with HISAT2 and HISAT-genotype. Nat Biotechnol 37, 907–915. doi: 10.1038/s41587-019-0201-4

Klein, K., Habiger, C., Iftner, T., and Stubenrauch, F. (2020). A TGF-β– and p63-Responsive Enhancer Regulates IFN-κ Expression in Human Keratinocytes. The Journal of Immunology. doi: 10.4049/jimmunol.1901178

Koster, M. I., Kim, S., Mills, A. A., DeMayo, F. J., and Roop, D. R. (2004). p63 is the molecular switch for initiation of an epithelial stratification program. Genes Dev. 18, 126–131. doi: 10.1101/gad.1165104

Kouwenhoven, E. N., Oti, M., Niehues, H., van Heeringen, S. J., Schalkwijk, J., Stunnenberg, H. G., et al. (2015). Transcription factor p63 bookmarks and regulates dynamic enhancers during epidermal differentiation. EMBO reports 16, 863–878. doi: 10.15252/embr.201439941

Krauskopf, K., Gebel, J., Kazemi, S., Tuppi, M., Löhr, F., Schäfer, B., et al. (2018). Regulation of the Activity in the p53 Family Depends on the Organization of the Transactivation Domain. Structure 26, 1091–1100.e4. doi: 10.1016/j.str.2018.05.013

Kulkarni, M. M., and Arnosti, D. N. (2003). Information display by transcriptional enhancers. Development 130, 6569–6575. doi: 10.1242/dev.00890

Lambert, S. A., Jolma, A., Campitelli, L. F., Das, P. K., Yin, Y., Albu, M., et al. (2018). The Human Transcription Factors. Cell 172, 650–665. doi: 10.1016/j.cell.2018.01.029

Lambourne, L., Mattioli, K., Santoso, C., Sheynkman, G., Inukai, S., Kaundal, B., et al. (2024). Widespread variation in molecular interactions and regulatory properties among transcription factor isoforms. bioRxiv, 2024.03.12.584681. doi: 10.1101/2024.03.12.584681

Latil, M., Nassar, D., Beck, B., Boumahdi, S., Wang, L., Brisebarre, A., et al. (2017). Cell-Type-Specific Chromatin States Differentially Prime Squamous Cell Carcinoma Tumor-Initiating Cells for Epithelial to Mesenchymal Transition. Cell Stem Cell 20, 191–204.e5. doi: 10.1016/j.stem.2016.10.018

LeBoeuf, M., Terrell, A., Trivedi, S., Sinha, S., Epstein, J. A., Olson, E. N., et al. (2010). Hdac1 and Hdac2 act redundantly to control p63 and p53 functions in epidermal progenitor cells. Dev Cell 19, 807–818. doi: 10.1016/j.devcel.2010.10.015

Lecellier, C.-H., Wasserman, W. W., and Mathelier, A. (2018). Human Enhancers Harboring Specific Sequence Composition, Activity, and Genome Organization Are Linked to the Immune Response. Genetics 209, 1055–1071. doi: 10.1534/genetics.118.301116

Lena, A. M., Rossi, V., Osterburg, S., Smirnov, A., Osterburg, C., Tuppi, M., et al. (2021). The p63 C-terminus is essential for murine oocyte integrity. Nat Commun 12, 383. doi: 10.1038/s41467-020-20669-0

Li, H., Handsaker, B., Wysoker, A., Fennell, T., Ruan, J., Homer, N., et al. (2009). The Sequence Alignment/Map format and SAMtools. Bioinformatics 25, 2078–2079. doi: 10.1093/bioinformatics/btp352

Li, L., Wang, Y., Torkelson, J. L., Shankar, G., Pattison, J. M., Zhen, H. H., et al. (2019). TFAP2C- and p63-Dependent Networks Sequentially Rearrange Chromatin Landscapes to Drive Human Epidermal Lineage Commitment. Cell Stem Cell 24, 271–284.e8. doi: 10.1016/j.stem.2018.12.012

Li, Y., Giovannini, S., Wang, T., Fang, J., Li, P., Shao, C., et al. (2023). p63: a crucial player in epithelial stemness regulation. Oncogene 42, 3371–3384. doi: 10.1038/s41388-023-02859-4

Lim, F., Solvason, J. J., Ryan, G. E., Le, S. H., Jindal, G. A., Steffen, P., et al. (2024). Affinity-optimizing enhancer variants disrupt development. Nature, 1–9. doi: 10.1038/s41586-023-06922-8

Lin-Shiao, E., Lan, Y., Welzenbach, J., Alexander, K. A., Zhang, Z., Knapp, M., et al. (2019). p63 establishes epithelial enhancers at critical craniofacial development genes. Science Advances 5, eaaw0946. doi: 10.1126/sciadv.aaw0946

Long, H. K., Prescott, S. L., and Wysocka, J. (2016). Ever-Changing Landscapes: Transcriptional Enhancers in Development and Evolution. Cell 167, 1170–1187. doi: 10.1016/j.cell.2016.09.018

Lopez-Pajares, V., Qu, K., Zhang, J., Webster, D. E., Barajas, B. C., Siprashvili, Z., et al. (2015). A LncRNA-MAF:MAFB Transcription Factor Network Regulates Epidermal Differentiation. Developmental Cell 32, 693–706. doi: 10.1016/j.devcel.2015.01.028

Marshall, C. B., Beeler, J. S., Lehmann, B. D., Gonzalez-Ericsson, P., Sanchez, V., Sanders, M. E., et al. (2021). Tissue-specific expression of p73 and p63 isoforms in human tissues. Cell Death Dis 12, 1–10. doi: 10.1038/s41419-021-04017-8

Massion, P. P., Taflan, P. M., Jamshedur Rahman, S. M., Yildiz, P., Shyr, Y., Edgerton, M. E., et al. (2003). Significance of p63 amplification and overexpression in lung cancer development and prognosis. Cancer Res 63, 7113–7121.

McDade, S. S., Henry, A. E., Pivato, G. P., Kozarewa, I., Mitsopoulos, C., Fenwick, K., et al. (2012). Genome-wide analysis of p63 binding sites identifies AP-2 factors as co-regulators of epidermal differentiation. Nucleic Acids Res 40, 7190–7206. doi: 10.1093/nar/gks389

McGrath, J. A., Duijf, P. H. G., Doetsch, V., Irvine, A. D., Waal, R. de, Vanmolkot, K. R. J., et al. (2001). Hay–Wells syndrome is caused by heterozygous missense mutations in the SAM domain of p63. Human Molecular Genetics 10, 221–230. doi: 10.1093/hmg/10.3.221

Melino, G., Memmi, E. M., Pelicci, P. G., and Bernassola, F. (2015). Maintaining epithelial stemness with p63. Sci. Signal. 8, re9–re9. doi: 10.1126/scisignal.aaa1033

Mills, A. A., Zheng, B., Wang, X. J., Vogel, H., Roop, D. R., and Bradley, A. (1999). p63 is a p53 homologue required for limb and epidermal morphogenesis. Nature 398, 708–13. doi: 10.1038/19531

Moore, J. E., Purcaro, M. J., Pratt, H. E., Epstein, C. B., Shoresh, N., Adrian, J., et al. (2020). Expanded encyclopaedias of DNA elements in the human and mouse genomes. Nature 583, 699–710. doi: 10.1038/s41586-020-2493-4

Moses, M. A., George, A. L., Sakakibara, N., Mahmood, K., Ponnamperuma, R. M., King, K. E., et al. (2019). Molecular Mechanisms of p63-Mediated Squamous Cancer Pathogenesis. Int J Mol Sci 20, 3590. doi: 10.3390/ijms20143590

Muerdter, F., Boryń, Ł. M., Woodfin, A. R., Neumayr, C., Rath, M., Zabidi, M. A., et al. (2018). Resolving systematic errors in widely used enhancer activity assays in human cells. Nat Methods 15, 141–149. doi: 10.1038/nmeth.4534

Murray-Zmijewski, F., Lane, D. P., and Bourdon, J.-C. (2006). p53/p63/p73 isoforms: an orchestra of isoforms to harmonise cell differentiation and response to stress. Cell Death Differ 13, 962–972. doi: 10.1038/sj.cdd.4401914

Napoli, M., Deshpande, A. A., Chakravarti, D., Rajapakshe, K., Gunaratne, P. H., Coarfa, C., et al. (2024). Genome-wide p63-Target Gene Analyses Reveal TAp63/NRF2-Dependent Oxidative Stress Responses. Cancer Research Communications 4, 264–278. doi: 10.1158/2767-9764.CRC-23-0358

Neumayr, C., Haberle, V., Serebreni, L., Karner, K., Hendy, O., Boija, A., et al. (2022). Differential cofactor dependencies define distinct types of human enhancers. Nature, 1–8. doi: 10.1038/s41586-022-04779-x

Ng, S. Y., Yoshida, N., Christie, A. L., Ghandi, M., Dharia, N. V., Dempster, J., et al. (2018). Targetable vulnerabilities in T- and NK-cell lymphomas identified through preclinical models. Nat Commun 9, 2024. doi: 10.1038/s41467-018-04356-9

Osterburg, C., and Dötsch, V. (2022). Structural diversity of p63 and p73 isoforms. Cell Death Differ 29, 921–937. doi: 10.1038/s41418-022-00975-4

Pastushenko, I., and Blanpain, C. (2019). EMT Transition States during Tumor Progression and Metastasis. Trends in Cell Biology 29, 212–226. doi: 10.1016/j.tcb.2018.12.001

Pattison, J. M., Melo, S. P., Piekos, S. N., Torkelson, J. L., Bashkirova, E., Mumbach, M. R., et al. (2018). Retinoic acid and BMP4 cooperate with p63 to alter chromatin dynamics during surface epithelial commitment. Nat Genet 50, 1658–1665. doi: 10.1038/s41588-018-0263-0

Peinado, H., Olmeda, D., and Cano, A. (2007). Snail, Zeb and bHLH factors in tumour progression: an alliance against the epithelial phenotype? Nat Rev Cancer 7, 415–428. doi: 10.1038/nrc2131

Peng, T., Zhai, Y., Atlasi, Y., ter Huurne, M., Marks, H., Stunnenberg, H. G., et al. (2020). STARR-seq identifies active, chromatin-masked, and dormant enhancers in pluripotent mouse embryonic stem cells. Genome Biology 21, 243. doi: 10.1186/s13059-020-02156-3

Perez, C. A., Ott, J., Mays, D. J., and Pietenpol, J. A. (2007). p63 consensus DNA-binding site: identification, analysis and application into a p63MH algorithm. Oncogene 26, 7363–7370. doi: 10.1038/sj.onc.1210561

Pickering, C. R., Zhou, J. H., Lee, J. J., Drummond, J. A., Peng, S. A., Saade, R. E., et al. (2014). Mutational landscape of aggressive cutaneous squamous cell carcinoma. Clin Cancer Res 20, 6582–92. doi: 10.1158/1078-0432.CCR-14-1768

Pokorna, Z., Vyslouzil, J., Vojtesek, B., and Coates, P. J. (2022). Identifying pathways regulating the oncogenic p53 family member ΔNp63 provides therapeutic avenues for squamous cell carcinoma. Cell Mol Biol Lett 27, 18. doi: 10.1186/s11658-022-00323-x

Qu, J., Tanis, S. E. J., Smits, J. P. H., Kouwenhoven, E. N., Oti, M., Van Den Bogaard, E. H., et al. (2018). Mutant p63 Affects Epidermal Cell Identity through Rewiring the Enhancer Landscape. Cell Reports 25, 3490–3503.e4. doi: 10.1016/j.celrep.2018.11.039

Qu, J., Yi, G., and Zhou, H. (2019). p63 cooperates with CTCF to modulate chromatin architecture in skin keratinocytes. Epigenetics Chromatin 12, 31. doi: 10.1186/s13072-019-0280-y

Quinlan, A. R., and Hall, I. M. (2010). BEDTools: a flexible suite of utilities for comparing genomic features. Bioinformatics 26, 841–842. doi: 10.1093/bioinformatics/btq033

Rahimov, F., Marazita, M. L., Visel, A., Cooper, M. E., Hitchler, M. J., Rubini, M., et al. (2008). Disruption of an AP-2alpha binding site in an IRF6 enhancer is associated with cleft lip. Nat. Genet. 40, 1341–1347. doi: 10.1038/ng.242

Ramírez, F., Ryan, D. P., Grüning, B., Bhardwaj, V., Kilpert, F., Richter, A. S., et al. (2016). deepTools2: a next generation web server for deep-sequencing data analysis. Nucleic Acids Res 44, W160–W165. doi: 10.1093/nar/gkw257

Ramsey, M. R., He, L., Forster, N., Ory, B., and Ellisen, L. W. (2011). Physical association of HDAC1 and HDAC2 with p63 mediates transcriptional repression and tumor maintenance in squamous cell carcinoma. Cancer Res 71, 4373–4379. doi: 10.1158/0008-5472.CAN-11-0046

Ramsey, M. R., Wilson, C., Ory, B., Rothenberg, S. M., Faquin, W., Mills, A. A., et al. (2013). FGFR2 signaling underlies p63 oncogenic function in squamous cell carcinoma. J Clin Invest 123, 3525–3538. doi: 10.1172/JCI68899

Ricci-Tam, C., Ben-Zion, I., Wang, J., Palme, J., Li, A., Savir, Y., et al. (2021). Decoupling transcription factor expression and activity enables dimmer switch gene regulation. Science 372, 292–295. doi: 10.1126/science.aba7582

Richardson, R., Mitchell, K., Hammond, N. L., Mollo, M. R., Kouwenhoven, E. N., Wyatt, N. D., et al. (2017). p63 exerts spatio-temporal control of palatal epithelial cell fate to prevent cleft palate. PLoS Genet. 13, e1006828. doi: 10.1371/journal.pgen.1006828

Riege, K., Kretzmer, H., Sahm, A., McDade, S. S., Hoffmann, S., and Fischer, M. (2020). Dissecting the DNA binding landscape and gene regulatory network of p63 and p53. eLife 9, e63266. doi: 10.7554/eLife.63266

Safieh, J., Chazan, A., Saleem, H., Vyas, P., Danin-Poleg, Y., Ron, D., et al. (2023). A molecular mechanism for the “digital” response of p53 to stress. Proceedings of the National Academy of Sciences 120, e2305713120. doi: 10.1073/pnas.2305713120

Sahu, B., Hartonen, T., Pihlajamaa, P., Wei, B., Dave, K., Zhu, F., et al. (2022). Sequence determinants of human gene regulatory elements. Nat Genet, 1–12. doi: 10.1038/s41588-021-01009-4

Saladi, S. V., Ross, K., Karaayvaz, M., Tata, P. R., Mou, H., Rajagopal, J., et al. (2017). ACTL6A Is Co-Amplified with p63 in Squamous Cell Carcinoma to Drive YAP Activation, Regenerative Proliferation, and Poor Prognosis. Cancer Cell 31, 35–49. doi: 10.1016/j.ccell.2016.12.001

Santos-Pereira, J. M., Gallardo-Fuentes, L., Neto, A., Acemel, R. D., and Tena, J. J. (2019). Pioneer and repressive functions of p63 during zebrafish embryonic ectoderm specification. Nat Commun 10, 3049. doi: 10.1038/s41467-019-11121-z

Senitzki, A., Safieh, J., Sharma, V., Golovenko, D., Danin-Poleg, Y., Inga, A., et al. (2021). The complex architecture of p53 binding sites. Nucleic Acids Research 49, 1364–1382. doi: 10.1093/nar/gkaa1283

Senoo, M., Pinto, F., Crum, C. P., and McKeon, F. (2007). p63 Is Essential for the Proliferative Potential of Stem Cells in Stratified Epithelia. Cell 129, 523–536. doi: 10.1016/j.cell.2007.02.045

Seo, J., Koçak, D. D., Bartelt, L. C., Williams, C. A., Barrera, A., Gersbach, C. A., et al. (2021). AP-1 subunits converge promiscuously at enhancers to potentiate transcription. Genome Res. 31, 538–550. doi: 10.1101/gr.267898.120

Sethi, I., Gluck, C., Zhou, H., Buck, M. J., and Sinha, S. (2017). Evolutionary re-wiring of p63 and the epigenomic regulatory landscape in keratinocytes and its potential implications on species-specific gene expression and phenotypes. Nucleic Acids Research 45, 8208–8224. doi: 10.1093/nar/gkx416

Sethi, I., Romano, R.-A., Gluck, C., Smalley, K., Vojtesek, B., Buck, M. J., et al. (2015). A global analysis of the complex landscape of isoforms and regulatory networks of p63 in human cells and tissues. BMC Genomics 16, 584. doi: 10.1186/s12864-015-1793-9

Sethi, I., Sinha, S., and Buck, M. J. (2014). Role of chromatin and transcriptional co-regulators in mediating p63-genome interactions in keratinocytes. BMC Genomics 15, 1042. doi: 10.1186/1471-2164-15-1042

Sheffield, N. C., Thurman, R. E., Song, L., Safi, A., Stamatoyannopoulos, J. A., Lenhard, B., et al. (2013). Patterns of regulatory activity across diverse human cell types predict tissue identity, transcription factor binding, and long-range interactions. Genome Res. 23, 777–788. doi: 10.1101/gr.152140.112

Slattery, M., Zhou, T., Yang, L., Dantas Machado, A. C., Gordân, R., and Rohs, R. (2014). Absence of a simple code: how transcription factors read the genome. Trends in Biochemical Sciences 39, 381–399. doi: 10.1016/j.tibs.2014.07.002

Song, E.-A. C., Min, S., Oyelakin, A., Smalley, K., Bard, J. E., Liao, L., et al. (2018). Genetic and scRNA-seq Analysis Reveals Distinct Cell Populations that Contribute to Salivary Gland Development and Maintenance. Sci Rep 8, 14043. doi: 10.1038/s41598-018-32343-z

Spitz, F., and Furlong, E. E. M. (2012). Transcription factors: from enhancer binding to developmental control. Nat Rev Genet 13, 613–626. doi: 10.1038/nrg3207

Strubel, A., Münick, P., Hartmann, O., Chaikuad, A., Dreier, B., Schaefer, J. V., et al. (2023). DARPins detect the formation of hetero-tetramers of p63 and p73 in epithelial tissues and in squamous cell carcinoma. Cell Death Dis 14, 1–12. doi: 10.1038/s41419-023-06213-0

Su, X., Paris, M., Gi, Y. J., Tsai, K. Y., Cho, M. S., Lin, Y.-L., et al. (2009). TAp63 prevents premature aging by promoting adult stem cell maintenance. Cell Stem Cell 5, 64–75. doi: 10.1016/j.stem.2009.04.003

Sundqvist, A., Vasilaki, E., Voytyuk, O., Bai, Y., Morikawa, M., Moustakas, A., et al. (2020). TGFβ and EGF signaling orchestrates the AP-1- and p63 transcriptional regulation of breast cancer invasiveness. Oncogene 39, 4436–4449. doi: 10.1038/s41388-020-1299-z

Szak, S. T., Mays, D., and Pietenpol, J. A. (2001). Kinetics of p53 Binding to Promoter Sites In Vivo. Molecular and Cellular Biology 21, 3375–3386. doi: 10.1128/MCB.21.10.3375-3386.2001

Taskiran, I. I., Spanier, K. I., Dickmänken, H., Kempynck, N., Pančíková, A., Ekşi, E. C., et al. (2024). Cell-type-directed design of synthetic enhancers. Nature 626, 212–220. doi: 10.1038/s41586-023-06936-2

Thomason, H. A., Zhou, H., Kouwenhoven, E. N., Dotto, G.-P., Restivo, G., Nguyen, B.-C., et al. (2010). Cooperation between the transcription factors p63 and IRF6 is essential to prevent cleft palate in mice. J Clin Invest 120, 1561–1569. doi: 10.1172/JCI40266

Thurfjell, N., Coates, P. J., Vojtesek, B., Benham-Motlagh, P., Eisold, M., and Nylander, K. (2005). Endogenous p63 acts as a survival factor for tumour cells of SCCHN origin. Int J Mol Med 16, 1065–1070.

Thurman, R. E., Rynes, E., Humbert, R., Vierstra, J., Maurano, M. T., Haugen, E., et al. (2012). The accessible chromatin landscape of the human genome. Nature 489, 75–82. doi: 10.1038/nature11232

Trauernicht, M., Martinez-Ara, M., and Steensel, B. van (2020). Deciphering Gene Regulation Using Massively Parallel Reporter Assays. Trends in Biochemical Sciences 45, 90–91. doi: 10.1016/j.tibs.2019.10.006

Trauernicht, M., Rastogi, C., Manzo, S. G., Bussemaker, H. J., and van Steensel, B. (2023). Optimisation of TP53 reporters by systematic dissection of synthetic TP53 response elements. Nucleic Acids Res 51, 9690–9702. doi: 10.1093/nar/gkad718

Truong, A. B., Kretz, M., Ridky, T. W., Kimmel, R., and Khavari, P. A. (2006). p63 regulates proliferation and differentiation of developmentally mature keratinocytes. Genes Dev. 20, 3185–3197. doi: 10.1101/gad.1463206

Tucci, P., Agostini, M., Grespi, F., Markert, E. K., Terrinoni, A., Vousden, K. H., et al. (2012). Loss of p63 and its microRNA-205 target results in enhanced cell migration and metastasis in prostate cancer. Proceedings of the National Academy of Sciences 109, 15312–15317. doi: 10.1073/pnas.1110977109

van Bokhoven, H., Hamel, B. C. J., Bamshad, M., Sangiorgi, E., Gurrieri, F., Duijf, P. H. G., et al. (2001). p63 Gene Mutations in EEC Syndrome, Limb-Mammary Syndrome, and Isolated Split Hand–Split Foot Malformation Suggest a Genotype-Phenotype Correlation. The American Journal of Human Genetics 69, 481–492. doi: 10.1086/323123

Verfaillie, A., Svetlichnyy, D., Imrichova, H., Davie, K., Fiers, M., Atak, Z. K., et al. (2016). Multiplex enhancer-reporter assays uncover unsophisticated TP53 enhancer logic. Genome Res. 26, 882–895. doi: 10.1101/gr.204149.116

Vu, H., and Ernst, J. (2022). Universal annotation of the human genome through integration of over a thousand epigenomic datasets. Genome Biology 23, 9. doi: 10.1186/s13059-021-02572-z

Wang, E. T., Sandberg, R., Luo, S., Khrebtukova, I., Zhang, L., Mayr, C., et al. (2008). Alternative isoform regulation in human tissue transcriptomes. Nature 456, 470–476. doi: 10.1038/nature07509

White, M. A., Myers, C. A., Corbo, J. C., and Cohen, B. A. (2013). Massively parallel in vivo enhancer assay reveals that highly local features determine the cis-regulatory function of ChIP-seq peaks. Proceedings of the National Academy of Sciences 110, 11952–11957. doi: 10.1073/pnas.1307449110

Wilson, V. G. (2013). “Growth and Differentiation of HaCaT Keratinocytes,” in Epidermal Cells, ed. K. Turksen (New York, NY: Springer New York), 33–41. doi: 10.1007/7651_2013_42

Woodstock, D. L., Sammons, M. A., and Fischer, M. (2021). p63 and p53: Collaborative Partners or Dueling Rivals? Frontiers in Cell and Developmental Biology 9. Available at: https://www.frontiersin.org/articles/10.3389/fcell.2021.701986 (Accessed September 14, 2022).

Wu, G., Yoshida, N., Liu, J., Zhang, X., Xiong, Y., Heavican-Foral, T. B., et al. (2023). TP63 fusions drive multicomplex enhancer rewiring, lymphomagenesis, and EZH2 dependence. Science Translational Medicine 15, eadi7244. doi: 10.1126/scitranslmed.adi7244

Yallowitz, A. R., Alexandrova, E. M., Talos, F., Xu, S., Marchenko, N. D., and Moll, U. M. (2014). p63 is a prosurvival factor in the adult mammary gland during post-lactational involution, affecting PI-MECs and ErbB2 tumorigenesis. Cell Death Differ 21, 645–654. doi: 10.1038/cdd.2013.199

Yanez-Cuna, J. O., Dinh, H. Q., Kvon, E. Z., Shlyueva, D., and Stark, A. (2012). Uncovering cis-regulatory sequence requirements for context-specific transcription factor binding. Genome Res 22, 2018–30. doi: 10.1101/gr.132811.111

Yang, A., Kaghad, M., Wang, Y., Gillett, E., Fleming, M. D., Dötsch, V., et al. (1998). p63, a p53 Homolog at 3q27–29, Encodes Multiple Products with Transactivating, Death-Inducing, and Dominant-Negative Activities. Molecular Cell 2, 305–316. doi: 10.1016/S1097-2765(00)80275-0

Yang, A., Zhu, Z., Kapranov, P., McKeon, F., Church, G. M., Gingeras, T. R., et al. (2006). Relationships between p63 Binding, DNA Sequence, Transcription Activity, and Biological Function in Human Cells. Molecular Cell 24, 593–602. doi: 10.1016/j.molcel.2006.10.018

Yoh, K. E., Regunath, K., Guzman, A., Lee, S.-M., Pfister, N. T., Akanni, O., et al. (2016). Repression of p63 and induction of EMT by mutant Ras in mammary epithelial cells. Proceedings of the National Academy of Sciences 113, E6107–E6116. doi: 10.1073/pnas.1613417113

Yu, X., Singh, P. K., Tabrejee, S., Sinha, S., and Buck, M. J. (2021). ΔNp63 is a pioneer factor that binds inaccessible chromatin and elicits chromatin remodeling. Epigenetics & Chromatin 14, 20. doi: 10.1186/s13072-021-00394-8

Zaret, K. S., and Mango, S. E. (2016). Pioneer transcription factors, chromatin dynamics, and cell fate control. Curr Opin Genet Dev 37, 76–81. doi: 10.1016/j.gde.2015.12.003

